# Programmed Cell Death-1 marks TOX^+^ T Regulatory Cells that Inhibit Autoimmunity and Anti-Tumor Immunity

**DOI:** 10.64898/2026.11.26.769460

**Authors:** Arda Akchengel, Jing Xuan Lim, Kathryn Sharples, Catherine Arden, Arian Laurence, Shoba Amarnath

## Abstract

Regulatory T cells (Tregs) are important for the maintenance of immune homeostasis. A proportion of peripheral Tregs constitutively express the coreceptor programmed cell death-1 (PD1), but the importance of this subset is yet to be investigated. We found PD1 marks TOX expressing Tregs. In both PD1 germline deficient and peripheral deficient Tregs, an increase in TOX^+^ Tregs were noted in steady state which was reproduced in antibody-based blocking studies. We next investigated if TOX^+^ Tregs were present in disease states. For this we utilised an experimental murine model of melanoma, colorectal cancer and autoimmune colitis. In all experimental conditions, TOX^+^ Tregs were present and marked by PD-1. To explore the function of PD1^+^TOX^+^ Tregs, we utilised a PD1-DTR switch. We found deleting PD1^+^ Tregs that marked TOX abrogated tumor suppression and enhanced T cell mediated colitis. Mechanistically, we found PD1^+^TOX^+^ Treg inhibited the functional plasticity of PD1^-^ Tregs. Taken together, this study identifies PD1 and TOX as markers of Treg function and highlights potential new immunotherapeutic approaches such as PD1^+^Treg cell depletion for enhanced anti-tumor responses.

## Introduction

Regulatory T cells (Tregs) defined by the master transcription factor FoxP3 are critical for maintaining immune homeostasis and preventing autoimmunity^1, 2^. In cancers, an increase in Tregs negatively impact tumor immunity. Depleting Tregs to enhance anti-tumor immunity has been widely investigated with most studies targeting the surface high-affinity receptor IL-2 (CD25) ^3, 4, 5^ which is constitutively expressed on Tregs.

In addition to CD25, Tregs also constitutively express several inhibitory co-receptors namely cytotoxic T-lymphocyte-associated protein 4 (CTLA-4), programmed cell death-1 receptor (PD1), T-cell immunoreceptor with Ig and ITIM domains^6, 7^ (TIGIT) etc. PD1 is a co-inhibitory receptor belonging to the B7-family and binds to its cognate ligands programmed death ligand-1 (PDL1) or programmed death ligand-2 (PDL2)^8^. PD1 when bound to PDL1 enhances FoxP3 expression in stimulated murine CD4^+^ T helper (Th) cells^9^, and maintains FoxP3 expression in human and murine Tbet^+^ Th1 cells^10^ ^11^. Within the tumor microenvironment (TME), PDL1 expression coincides with increased intra-tumoral FoxP3^+^ T cells suggesting a role for this signaling pathway in contributing towards the maintenance of FoxP3^+^ T cells within the TME.

Blocking antibodies to CTLA4 and PD1 has shown tremendous success in driving tumor immunity, but a majority of the cancer population develop resistance towards these therapies, partly due to enhanced Treg function within tumors. Peripheral deletion of PD1 in Tregs in experimental murine models of autoimmunity^12^, infectious disease^13^ and cancer^14^has identified PD1 as an inhibitor of Treg function. Blocking antibodies to PD1 unleash this inhibitory signal, whereby resistance to tumors develop through selective expansion of suppressive Tregs^15^. Indeed, peripheral deletion of PD1 in Tregs drives the expansion of several functional differentiation states within the tumor microenvironment (TME), with an evolution of a significantly suppressive Treg population that is marked by tumor necrosis factor family of receptors^14^. In addition to the suppressive state, PD1 deficiency also drove a type 1 phenotype in Tregs and a cytotoxic phenotype^14^. These reports suggest that PD1 has the capacity to remodel Treg functional states. While the intrinsic importance of PD1 in Treg stability has been extensively explored, to date the phenotype of naturally occurring PD1^+^ Tregs and their function is yet to be defined.

A subset of Tregs constitutively express PD1 which have not been fully characterized and given the role of PD1 in driving Treg differentiation states, it remains possible that this subset could be important in driving peripheral Treg function. Although blocking antibodies to PD1 has been shown to enhance Treg numbers, to date there are no reports on deleting PD1^+^ Tregs and their role remains unresolved in Treg biology.

In CD8^+^ T cell, PD1 has been shown to inhibit exhaustion^16^, a process that is sustained by the master transcription factor TOX, which is expressed in both progenitor of exhausted (Tpex) and exhausted (Tex) CD8^+^ T cells^17^. TOX is a high-mobility-group (HMG) box transcription factor with key function in driving peripheral CD8^+^ T cells. TOX belongs to a larger family of transcription factors (TF) including TOX, TOX2, TOX3 and TOX4 and enables DNA recognition and binding that is structure dependent and sequence independent^18, 19^. TOX and TOX2 are expressed in hematopoietic cells populations^18^. TOX plays a key role in the development of CD4^+^ thymocytes^19, 20^, natural killer (NK) cell development and in the differentiation of common lymphoid progenitors to innate lymphoid cells by controlling ThPOK. In the periphery^21^, TOX is upregulated following antigen/TCR stimulation and/or inflammatory cytokines in CD8^+^ T cells^22^. In tumors^23^ and chronic viral infections^24^, TOX expression progressively increases with time, driven by continuous antigen encounter and TCR stimulation. Forced expression of TOX in CD8^+^ T cells result in a Tex phenotype with reduced cytokine expression and enhanced expression of PD1 and TCF1^17^. In early chronic viral infections, the expression of TOX is dependent on antigen dose and is corelated with PD1 expression in CD8^+^ T cells^24^. By regulating inhibitory receptor expression such as PD1^17, 25^, TOX can control overstimulation of CD8^+^ T cell survival and persistence. TOX deficiency results in the loss of inhibitory co-receptor expression, but it does not control effector function or memory formation, and CD8^+^ T cells can transition into dysfunctional states suggesting TOX is important for persistence and survival but does not control effector function^17, 23, 24^. In peripheral CD4^+^ T cells, TOX has been shown to be critical for Tfh cell differentiation^26,27^ during acute infection and overexpression drives IL10 productionin CD4^+^ T cells *in vitro*^28^. Recently, the role of TOX in effector CD4^+^ T cells has been further investigated where it has been identified as a key driver of Th1 cell differentiation^29^. TOX expression in CD4^+^ T cells in cancer was associated with increased cytotoxicity, anti-tumor immunity and improved response to immunotherapy. TOX expression has been reported in Tregs in human malignancies including non-Hodgkin lymphoma^30^, and multiple myeloma^31^. TOX is also induced during tonic signaling in CAR-engineered human Treg cells^32^. Recently, TOX expression in tumor derived Tregs has been reported whereby it drives an effector like Treg state^33^ highlighting its importance in Treg fitness and stability.

In the current work, we explored the link between PD1 and TOX expression in Tregs and its role in Treg function. We found that PD1 marked TOX expression in peripheral Tregs. Both germline deletion, peripheral deletion or anti-PD1 antibody treatment resulted in expansion of the TOX^+^ Treg subset in steady state. These data suggest that PD1 both marks and restrains TOX^+^Tregs. The PD1^+^TOX^+^ subset was increased within the TME and decreased in inflammatory dextran sodium sulfate (DSS) driven colitis. Deletion of PD1^+^Tregs using a diphtheria toxin switch enhanced anti-tumor immunity and exacerbated experimental colitis. These data are the first to define a new modality for overcome checkpoint inhibitor resistance through, deletion of PD1^+^Tregs to overcome anti-PD1 mediated resistance in cancers.

## Results

### PD1 marks TOX^+^ Tregs in steady state in both secondary lymphoid organs and healthy tissues

The expression of TOX is associated with early chronic viral infections showing a dependency on antigen dose and is corelated with PD1 expression in CD8^+^ T cells^24^. This observation led us to investigate the corelation of PD1 to TOX in Tregs. Flow cytometry analysis of splenocytes showed a direct protein co-expression between PD1 and TOX in Tregs (**Extended Fig. 1a)**. We next investigated the expression of TCF1 within this population, given the relationship between TCF1 and TOX in driving CD8^+^ T cell differentiation from Tpex to Tex^34^. We found that PD1 marked both TCF1^+^TOX^+^ and TCF1^-^ TOX^+^ population in splenocytes of healthy mice (**Fig. 1a-b**). We next sub-divided the PD1^+^Tregs into negative, medium and high expressors to determine which subset had the highest TCF1 and TOX expression (**Extended Fig. 1b**). Flow cytometry analysis demonstrated that PD1^hi^Tregs were significantly enriched in both TCF1 and TOX populations (**Extended Fig.1c**). On further phenotyping, we found that PD1^+^TCF1^-^ TOX^+^ Tregs had significantly decreased BCL6 expression, no change in CD73, had significantly elevated KLRG1 expression and no change in CD127 with a significant decrease in CD44^+^CD62L^+^ phenotype and an increase in CD44^+^CD62L^-^ phenotype compared to the PD1^+^ TCF1^+^TOX^+^ Tregs (**Fig. 1c-i**). These data suggest that PD1 marks a population of TOX^+^ Tregs, and within the TCF1^-^TOX^+^ subset, a decrease in central memory phenotype but an activated effector phenotype was seen. We next investigated whether PD1 marks TCF1^+^TOX^+^ and TCF1^-^ TOX^+^populations in secondary lymphoid organs, and tissues. A similar profile was noted in mesenteric lymph nodes (MLN), liver, lung, lamina propria lymphocytes (LPLs) in the small intestine and colon (**Fig. 1j-o**). Next, PD1^+^Tregs were divided into medium and high expressors. Again, PD1^hi^Tregs significantly marked TCF1^+^TOX^+^ Tregs and TCF1^-^TOX^+^ Tregs (**Extended Fig.1d-h**) in all tissues tested. Further phenotyping of both populations in the various organs showed a similar profile to splenocytes whereby either a decrease or a trend towards decreased BCL6 and an increase in KLRG1 was noted in the TCF1^-^TOX^+^ Treg subset (**Extended Fig. 2a-e**). Of note no difference in CD44^+^CD62L^-^ expression was noted in all the tissues tested unlike splenocytes suggesting a lack of activation of this subset in tissues. We further interrogated the presence of these cells in the circulation namely blood and in other tissues such as kidneys and pancreas by analyzing a publicly mined murine scRNA-seq dataset ^35^. Here we found that both populations were present in the LPL, pancreas and kidney but minimal in the blood and liver (**Extended Fig.3a-e; Supplementary Table 1**). To determine the differentiation trajectory of the TCF1^+^TOX^+^Tregs, a psuedotime trajectory analysis of the dataset was performed. In all three tissues, TCF1^+^TOX^+^ population led to the TCF1^-^TOX^+^ population followed by differentiating into other phenotypes (Pancreas lineage 1-3; LPL lineage 3; Kidneys lineage3. (**Extended Fig. 3&4**). Hence, like CD8^+^ T cell differentiation, we suggest that PD1 marks a population of Tregs that express TCF1^+^TOX^+^ transcription factors and this population can differentiate into TCF1^-^TOX^+^ Tregs.

**Figure 1:**
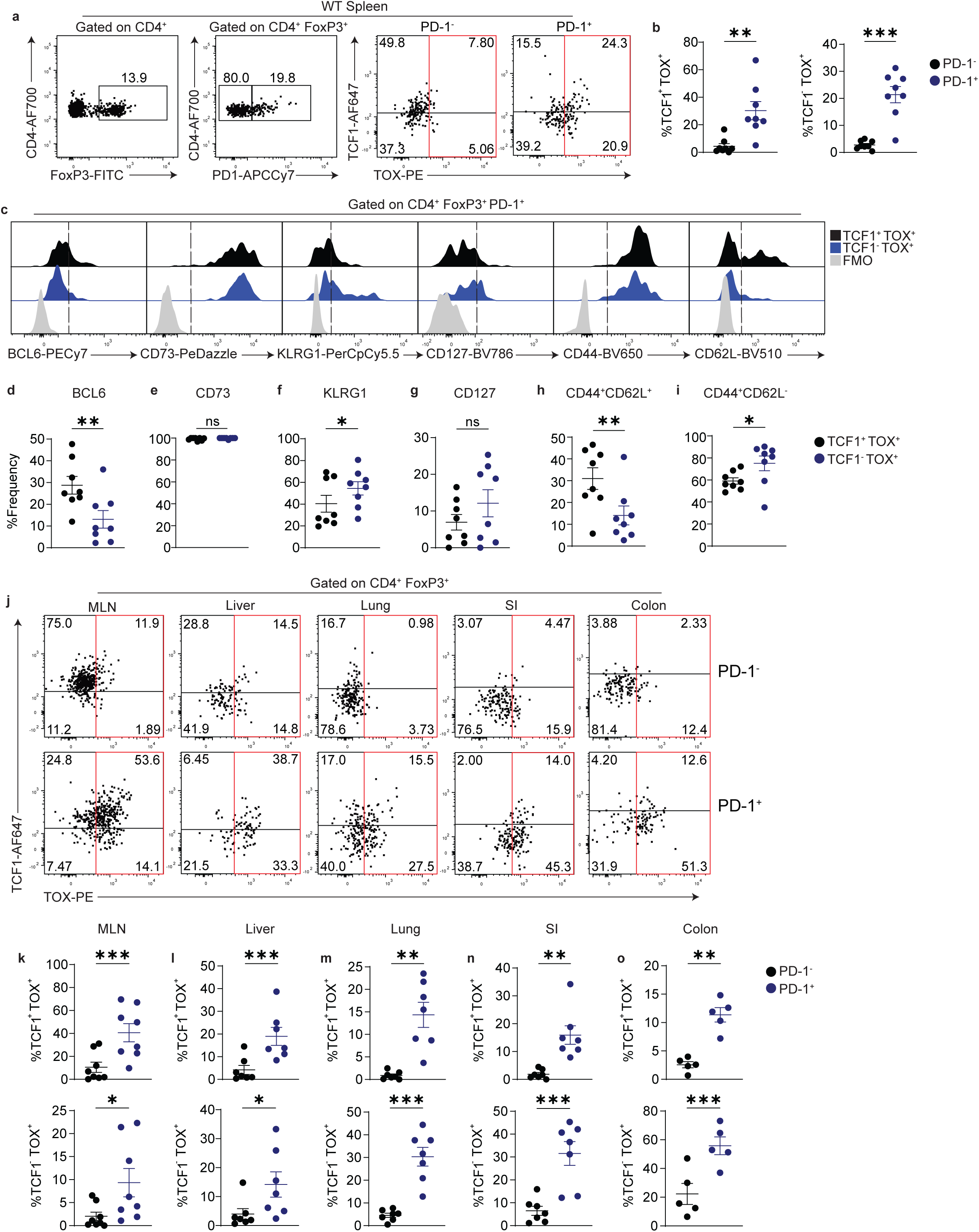
PD1 marks TOX^+^ T regulatory cells in wildtype (WT) mice. C57BL6*Foxp3*^RFP^ mice splenocytes were characterised using flow cytometry for TOX^+^ Treg subsets. Splenocytes were gated on CD4 and FoxP3, followed by gating on PD1^neg^ and PD1^pos^ subsets. From this subset TCF1 and TOX analysis was performed and representative flow cytometry shown (**a**). Summary of the frequency of TCF1^+^TOX^+^and TCF1^-^TOX^+^ cells (**b**) Further characterisation of TCF1^+^TOX^+^ cells and TCF1^-^TOX^+^ cells were performed for proteins namely BCL6, CD73, KLRG1, CD127, CD44 and CD62L. Representative flow plot for all markers along with FMOs (**c**), summary of the frequency of BCL6 (**d**), CD73 (**e**), KLRG1 (**f**), CD127 (**g**), CD44^+^CD62L^+^ (**h**), CD44^+^CD62L^-^ (**i**). Tregs from mesenteric lymph nodes (MLN), liver, lung, small intestine LPL and colonic LPL were further characterised for TCF1 and TOX in PD1^-^ and PD1^+^ subsets. Representative flow plot of TCF1 and TOX expression in various tissues (**j**). Frequency of TCF1^+^TOX^+^ cells and TCF1^-^TOX^+^ within CD4^+^FoxP3^+^PD1^-^ and CD4^+^FoxP3^+^PD1^+^ Tregs in MLN (**k**), liver (**l**), lung (**m**), small intestine (**n**) and colonic LPLs (**o**). Each dot represents a mouse, data shown are Mean±SEM, a paired student t test was performed. \**P* ≤ 0.05, \*\**P* ≤ 0.01, \*\*\**P* ≤ 0.001, \*\*\*\**P* ≤ 0.0001. FMO, fluorescence-minus-one control.

### Germline PD1 deficiency increases TOX^+^ Tregs in steady state

We next investigated if PD1 was merely a marker for TOX^+^ Tregs or whether it had an intrinsic regulatory role in this population. B6.WT and *Pdcd1^-/-^* mice secondary lymphoid organs and tissues were phenotype by flow cytometry. In the spleen, no changes were noted both in frequency and absolute counts of TCF1^+^TOX^+^ cells, but a significant increase in both frequency and numbers were noted in the TCF1^-^TOX^+^ Tregs (**Fig. 2a-c**). No change in the frequency and number of CD73, KLRG1 was noted in the TCF1^+^TOX^+^ between WT and *Pdcd1^-/-^*mice, but a significant upregulation in the frequency of CD44^+^CD62L^-^ cells were noted (**Fig. 2d-h**). Within the TCF1^-^TOX^+^ population, we found a significant increase in the absolute numbers of CD73, KLRG1 and CD44^+^CD62L^-^ cells. These data suggest that PD1 deficiency selectively increased TCF1^-^TOX^+^ Tregs which were characterized by an activation profile (**Fig. 2i-m**). When MLNs and other tissues were interrogated, the phenotype was protected whereby the frequency and number of TOX^+^ Tregs were significantly increased (**Extended Fig.5**). These data suggest that PD1 directly controls TOX^+^ subset in steady state.

**Figure 2:**
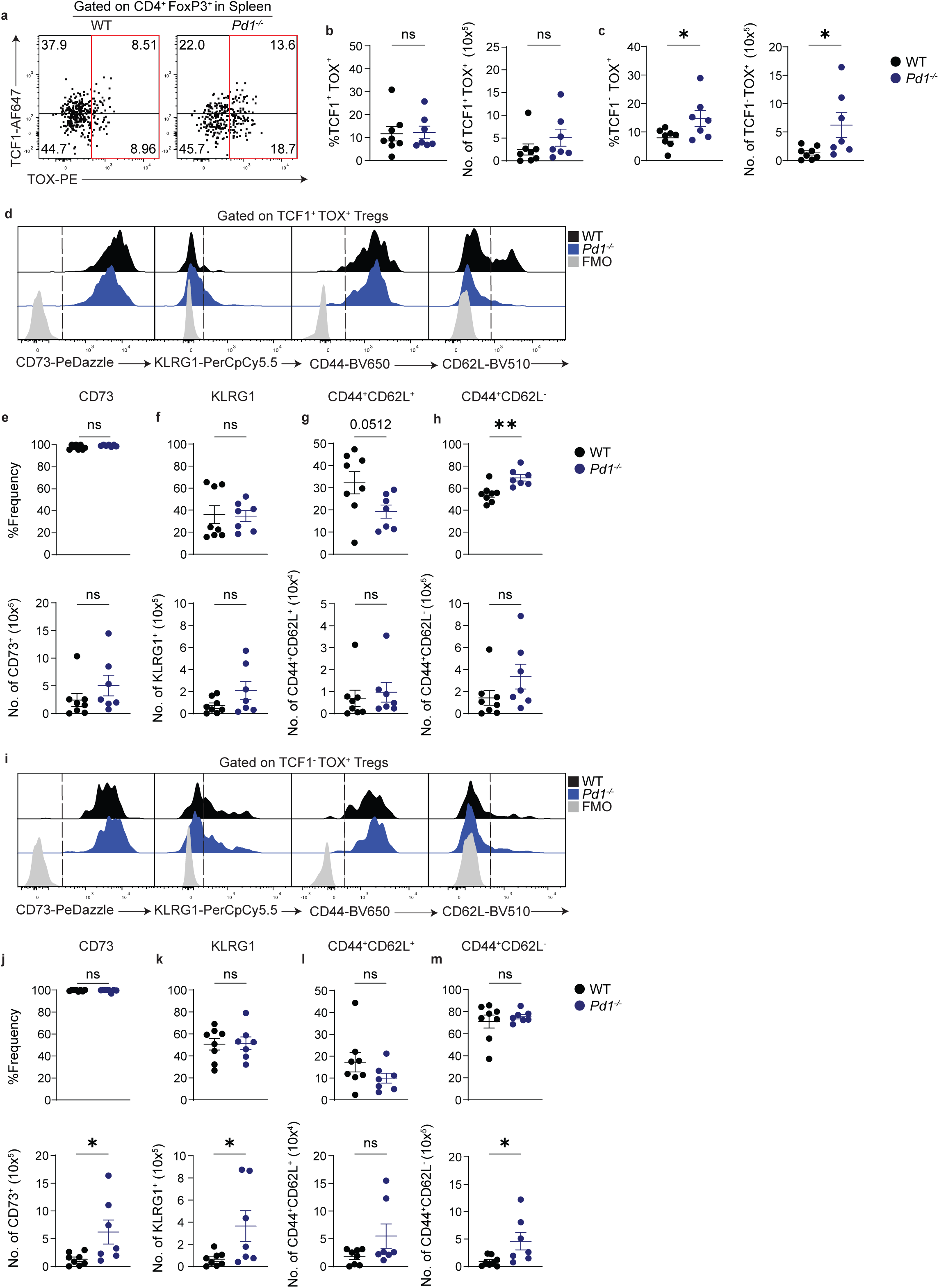
Germline PD1 deficiency increases TOX^+^ T regulatory cells. C57BL6 *Foxp3^RFP^* and *Pdcd1^-/-^Foxp3^RFP^* mice splenocytes were characterised using flow cytometry for TOX^+^ T regulatory subsets. Representative flow plot of TCF1 and TOX from Tregs (**a**), frequency and absolute numbers of TCF1^+^TOX^+^ subset (**b**), frequency and absolute numbers of TCF1^-^TOX^+^ subset (**c**). Further characterisation of TCF1^+^TOX^+^ subset was performed using markers such as CD73, KLRG1, CD44 and CD62L. Representative flow plot of marker expression (**d**), frequency and absolute counts of CD73 (**e**), KLRG1 (**f**), CD44^+^CD62L^+^(**g**), CD44^+^CD62L^-^ (**h**). Further characterisation of TCF1^-^TOX^+^ subset was performed using markers such as CD73, KLRG1, CD44 and CD62L. Representative flow plot of marker expression (**i**), frequency and absolute counts of CD73 (**j**), KLRG1 (**k**), CD44^+^CD62L^+^ (**l**), CD44^+^CD62L^-^ (**m)**. Each dot represents a mouse, data shown are Mean±SEM, a un-paired student t test was performed. \**P* ≤ 0.05, \*\**P* ≤ 0.01, \*\*\**P* ≤ 0.001, \*\*\*\**P* ≤ 0.0001. FMO, fluorescence-minus-one control.

### Peripheral deletion or antibody mediated blocking of PD1 increases TCF1^-^TOX^+^ Tregs in steady state

Since germline deletion of PD1 can result in cumulative accumulation of Tregs, we next investigated if peripheral deletion which mirror anti-PD1 treatment can enhance both TCF1^+^TOX^+^ and TCF1^-^TOX^+^ subsets. For this analysis, *Foxp3^Cre-ERT2^Pd1^flfl-DTR-tdtomato^*mice were either treated with PBS or tamoxifen for 5 days and then spleen, MLN and other tissues were analyzed using flow cytometry. In this scenario, a decrease in the frequency of TCF1^+^TOX^+^ Treg subset were noted in the splenocytes on tamoxifen treatment (**Fig. 3a-b**) and an increase in TCF1^-^TOX^+^ subset was noted both with respect to frequency and cell number (**Fig. 3a&c**). Further characterization of the TCF1^+^TOX^+^ Treg subset showed no difference in CD73, KLRG1 or CD44^+^CD62L^-^ population both with respect to frequency and absolute counts (**Fig. 3d-h**). Within the TCF1^-^TOX^+^ Treg subset, although no change in frequency was noted, we found a significant increase in CD73, KLRG1 and CD44^+^CD62L^-^ absolute counts (**Fig. 3i-m**). On phenotyping MLNs, no change in frequency or absolute counts were noted within the TCF1^+^TOX^+^ Treg subsets but we found an increase in absolute numbers in the TCF1^-^TOX^+^ Treg subset (**Extended Fig. 6a**). Within tissue, a decrease in the frequency was noted in the TCF1^+^TOX^+^ Treg subsets but an increase in absolute numbers was found in the TCF1^-^TOX^+^ Treg subset (**Extended Fig. 6b**). Next, we investigated if blocking PD1 using an antibody will generate a similar phenotype. We found that there was no change in the frequency and absolute numbers of TCF1^+^TOX^+^ Treg subsets but a significant increase in both frequency and absolute counts of TCF1^-^TOX^+^ Treg subset was seen (**Extended Fig. 7a-c**). Further characterization of TCF1^+^TOX^+^ subset showed no difference in CD73, KLRG1 or CD44^+^CD62L^-^ Treg population both with respect to frequency and absolute counts (**Extended Fig. 7d-e**). Within the TCF1^-^TOX^+^ Treg subset, a significant increase in frequency and absolute counts of CD73, and CD44^+^CD62L^-^ were observed (**Extended Fig. 7f-g**). These data suggest that PD1 specifically regulates the generation of TCF1^-^TOX^+^ subset with an effector Treg phenotype.

**Figure 3:**
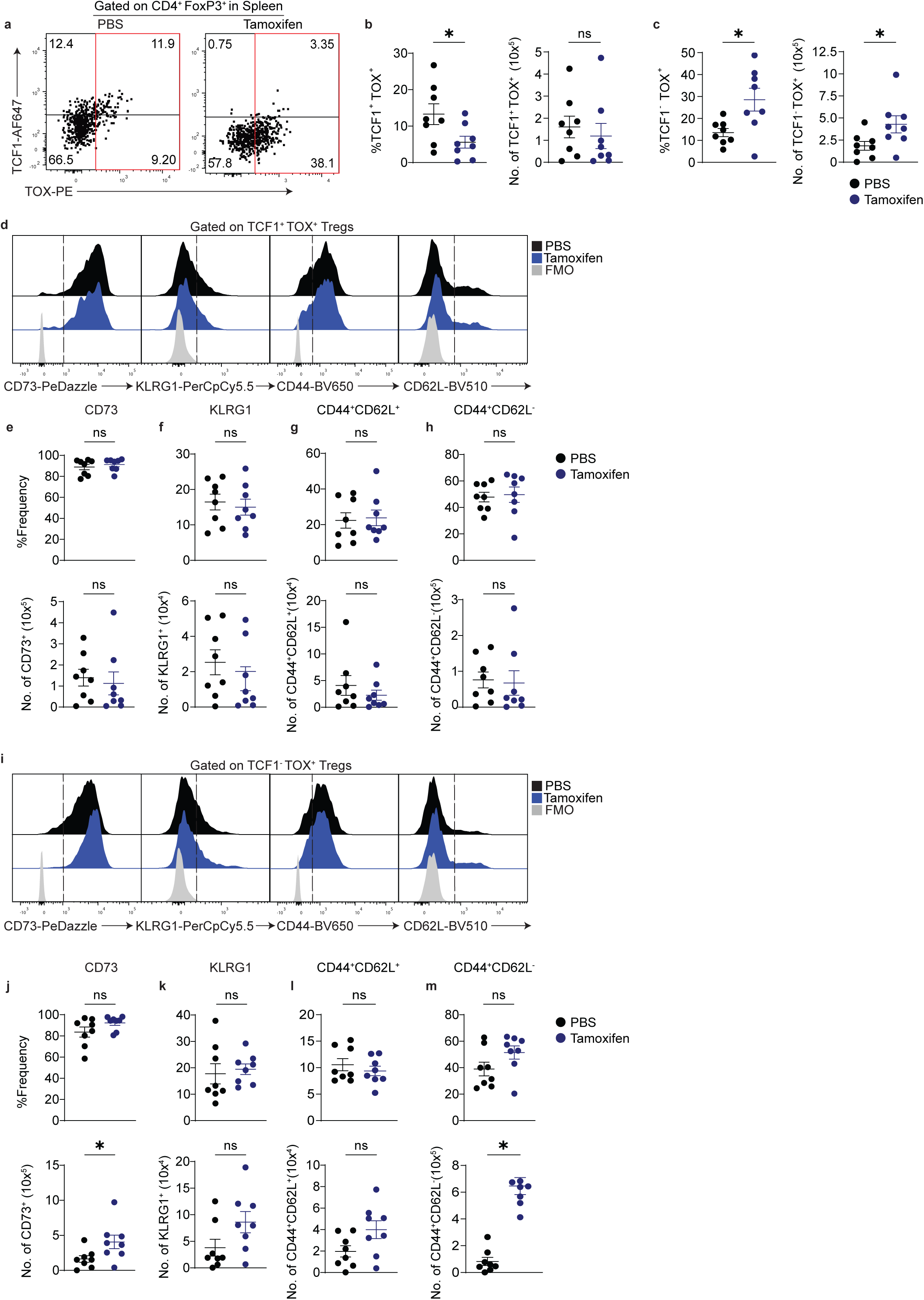
Peripheral PD1 deficiency increases TOX^+^ T regulatory cells. B6. *Pdcd ^fl/fl-DTR-Tdtomato^Foxp3^cre-ERT2^*mice were treated with either PBS or tamoxifen for 5 days and then splenocytes were harvested for flow cytometry characterisation. Representative flow plot of TCF1 and TOX from T regulatory cells (**a**), frequency and absolute numbers of TCF1^+^TOX^+^ subset (**b**), frequency and absolute numbers of TCF1^-^TOX^+^ subset (**c**). Further characterisation of TCF1^+^TOX^+^ subset was performed using markers such as CD73, KLRG1, CD44 and CD62L. Representative flow plot of marker expression (**d**), frequency and absolute counts of CD73 (**e**), KLRG1 (**f**), CD44^+^CD62L^+^(**g**), CD44^+^CD62L^-^ (**h**). Further characterisation of TCF1^-^TOX^+^ subset was performed using markers such as CD73, KLRG1, CD44 and CD62L. Representative flow plot of marker expression (**i**), frequency and absolute counts of CD73 (**j**), KLRG1 (**k**), CD44^+^CD62L^+^ (**l**), CD44^+^CD62L^-^ (**m)**. Each dot represents a mouse, data shown are Mean±SEM, an un-paired student t test was performed. \**P* ≤ 0.05, \*\**P* ≤ 0.01, \*\*\**P* ≤ 0.001, \*\*\*\**P* ≤ 0.0001. FMO, fluorescence-minus-one control.

### PD1^+^TOX^+^ Tregs are increased in B16F10 Melanoma and MC38 colorectal cancer tumor microenvironment

B16F10 melanoma cells were inoculated into *Pd1^fl/fl-DTR-Tomato^* mice and tumors were allowed to engraft and grow (**Extended Fig. 8a**). For TOX^+^ Tregs to have a functional impact, we hypothesized that a substantial expansion of PD1^+^ Tregs should occur within the TME. In line with this hypothesis, we found a significant increase in PD1^+^ Tregs in the TME, tumor draining lymph node (tdLN), and spleen (**Extended Fig. 8b-d**). On further analysis, it was found that similar to the steady state, PD1 specifically marked both TCF1^+^TOX^+^ and TCF1^-^ TOX^+^ subsets in the TME, tdLN and spleen. In addition, a significant increase in the frequency of these subsets were also noted in tumor bearing mice (**Fig. 4a-c**). Similar to steady state data, TCF1^-^TOX^+^ Tregs significantly downregulated BCL6, but upregulated KLRG1. Furthermore, we measured the expression of TIM3 which can be upregulated in CD8^+^ Tex cells in the TME. Indeed, an increase in TIM3 was noted within the TCF1^-^TOX^+^ Tregs in the TME (**Fig. 4d-g**). These data suggest that the TCF1^-^TOX^+^ Treg subsets significantly increase in tumor and may have an impact on tumor immunotherapy. To further confirm these findings, we used the MC38 colorectal cancer tumor model. Mice were inoculated with MC38 and then tumors and spleen were harvested for immunobiology. An increase in PD1^+^Tregs within the TME was again noted (**Extended Fig. 8e)** but unlike melanoma a significant increase in TCF1^-^TOX^+^ Treg subset was noted both in TME and spleen, but an increase in TCF1^+^TOX^+^ Treg was found only in the spleen (**Fig. 4h-j**). Similar to melanoma, TCF1^-^TOX^+^ Treg subsets did not express BCL6, showed no change in TIM3 and had elevated KLRG1 expression (**Fig. 4k-m**). Consolidating data from both TME, we can conclude that PD1^+^ Tregs are significantly increased in the TME, and mark TCF1^-^TOX^+^ subset which express the activation marker KLRG1.

**Figure 4:**
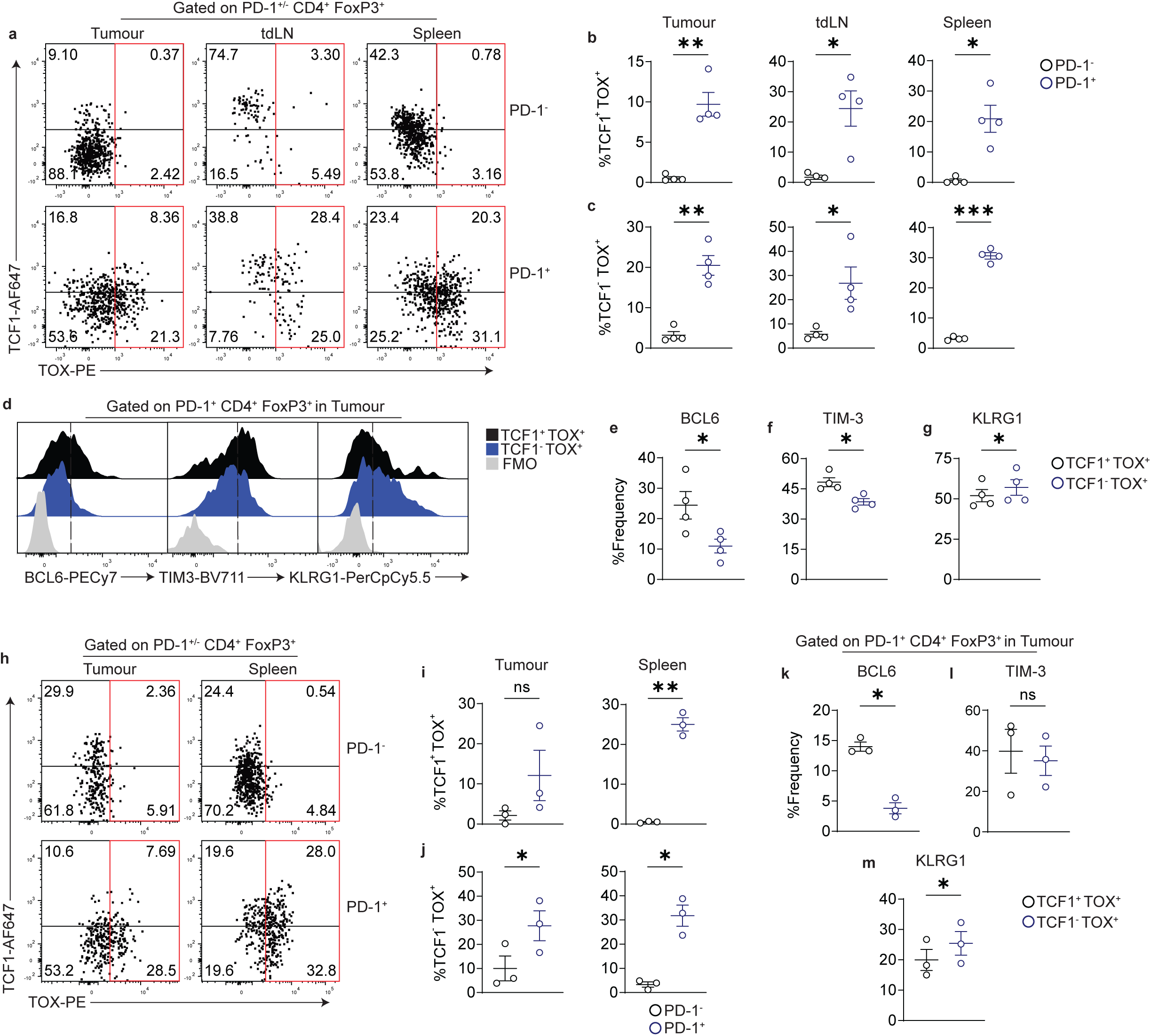
TOX^+^ Tregs are marked by PD1 within the B16F10 and MC38 TME. Tumors were harvested and flow cytometry characterisation of tumor derived lymphocytes (TILs), tumor draining lymph nodes (tdLN) and splenocytes was performed. Representative flow plot of TCF1 and TOX within the PD1^-^FoxP3^+^ Tregs and PD1^+^FoxP3^+^ Tregs in TILs, tdLN and Spleen (**a**), Summary of the frequency of TCF1^+^TOX^+^ Tregs in TILs, tdLN and spleen within PD1^+^ and PD1^-^ Treg subsets (**b**), summary of the frequency of TCF1^-^TOX^+^ Tregs in TILs, tdLN and spleen within PD1^+^ and PD1^-^ Treg subsets (**c**). TCF1^+^TOX^+^ and TCF1^-^TOX^+^ Tregs were further characterised for BCL6 expression, TIM3 and KLRG1. Representative flow cytometry profile is shown in (**d**), Summary data of BCL6 expression (**e**), TIM-3 (**f**), and KLRG1 (**g**). Mice were inoculated with 2×10^5^ MC38 cells and then TILs were characterised. Representative flow plot showing TCF1 versus TOX in PD1^+^ and PD1^-^ Treg subsets in TILs and spleen (**h**). Cumulative data showing the frequency of TCF1^+^TOX^+^ (**i**) and TCF1^-^TOX^+^ cells in PD1^+^ and PD1^-^ Treg subsets (**j**). Cumulative data on BCL6 (**k**), TIM3 (**l**) and KLRG1 (**m**) in TCF1^+^TOX^+^ and TCF1^-^TOX^+^ subsets is shown. Each dot represents a mouse, data shown are Mean±SEM, a paired student t test was performed. \**P* ≤ 0.05, \*\**P* ≤ 0.01, \*\*\**P* ≤ 0.001, \*\*\*\**P* ≤ 0.0001.

**Figure 5:**
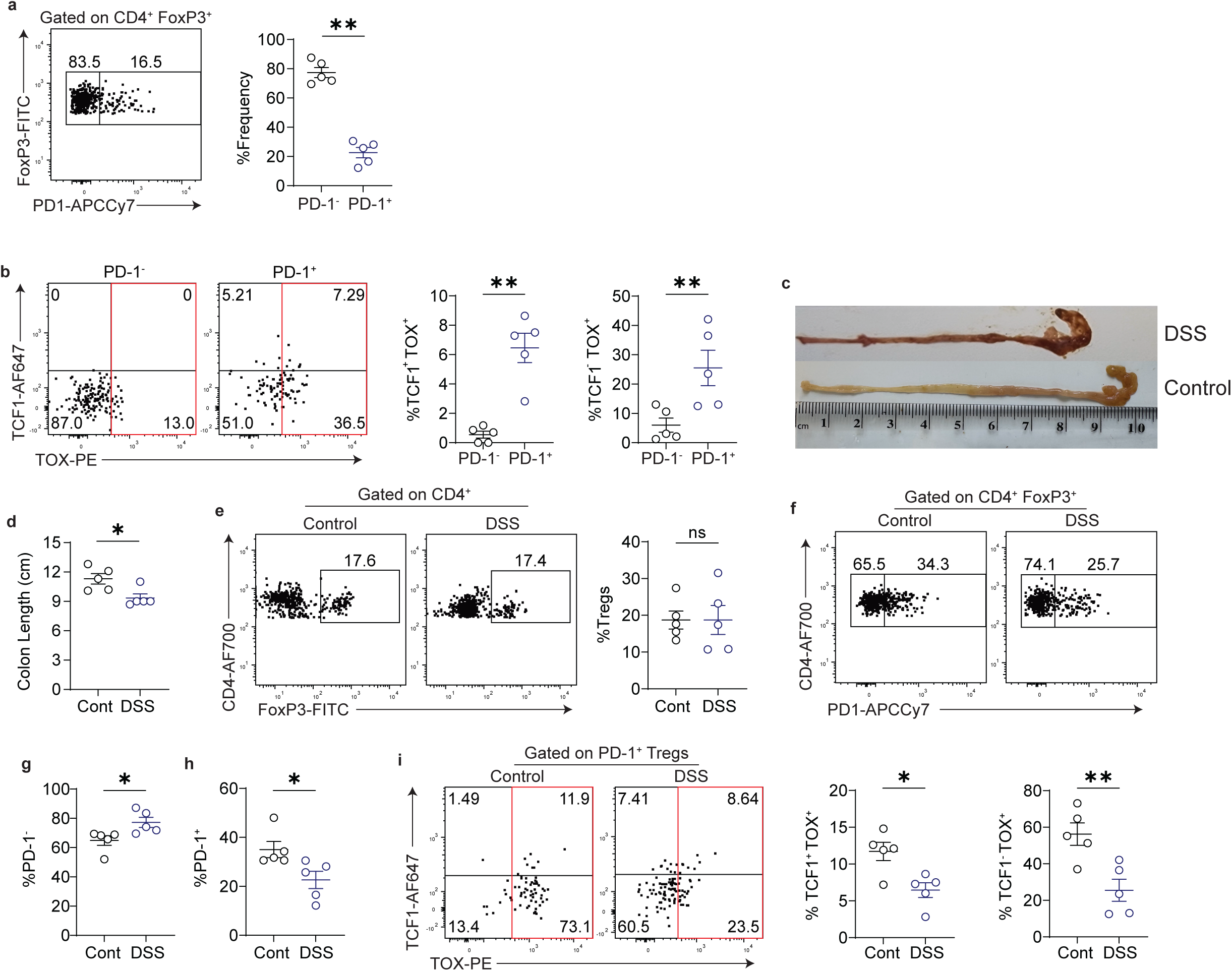
PD1^+^TOX^+^ Tregs are decreased in DSS colitis compared to healthy colons. *Foxp3^RFP^* mice were treated with DSS in water and then Tregs were characterised in the LPLs. Representative flow plot showing FoxP3 expression and PD1 expression and frequency of PD1^-^ and PD1^+^ Tregs (**a**). Representative flow plot of TCF1 and TOX within PD1^+^ Tregs (**b**) and cumulative frequency of TCF1^+^TOX^+^ and TCF1^-^TOX^+^ subsets in LPL. DSS treated mice were compared with WT mice to determine changes in PD1^+^ Tregs and TOX^+^ Tregs in steady state and inflammation. Colon length was measured post DSS (**c-d**), Frequency of FoxP3^+^ cells in WT and DSS mice (**e**), frequency of PD1^+^Tregs and PD1^-^Tregs in WT and DSS mice (**f-h**), Frequency of TCF1 and TOX subsets in WT and DSS mice (**i**). Each dot represents a mouse, data shown are Mean±SEM, a paired (**a-b**) and un-paired (**d-i**) student t test was performed. \**P* ≤ 0.05, \*\**P* ≤ 0.01, \*\*\**P* ≤ 0.001, \*\*\*\**P* ≤ 0.0001.

### PD1^+^TOX^+^ Tregs are decreased in DSS colitis compared to healthy colons

To determine if the increase in PD1^+^Tregs and the subsequent increase in TOX^+^ Tregs is restricted to the TME, we tested the phenomenon in a murine model of dextran sodium sulphate (DSS) mediated colitis. In contrast to the TME, a decrease in the frequency of PD1^+^ Tregs was noted in the colonic LPL (**Fig. 5a**). However, both TCF1^+^TOX^+^ and TCF1^-^TOX^+^ Treg subsets were significantly restricted to the PD1^+^ Treg compartment within the LPLs (**Fig. 5b**). We next compared the frequency of these subsets with normal LPL. DSS treatment resulted in the decrease in the colon length as expected (**Fig. 5c-d**). There was no change in the frequency of Tregs was noted when compared to WT mice (**Fig. 5e**), but a significant decrease in PD1^+^Tregs were found within the LPLs (**Fig. 5f-h**). In line with this decrease, we found a significant decrease in both TCF1^+^TOX^+^ and TCF1^-^TOX^+^ Treg subsets (**Fig. 5i**). These data suggest that in inflammation, there is a significant decrease in PD1^+^Tregs within the LPLs and this marks a significant downregulation of TOX^+^ Treg subsets.

### Deleting PD1^+^ Tregs with a diphtheria toxin switch abrogates protection against colitis and enhances B16 F10 melanoma tumor Immunity

Since PD1 marked both TCF1^+^TOX^+^ and TCF1^-^TOX^+^ Tregs, deleting PD1^+^ cells can provide direct evidence on the function of TOX^+^ Tregs. For these experiments, we utilised the *PD1^fl/fl-DTR-Tdtomato^* whereby splenocytes were cultured in the presence of diphtheria toxin (DT) and the depletion of PD1 expressing T cells was measured by flow cytometry (**Extended Fig. 9a-e**). On optimizing DT concentration and validating deletion of PD1^+^ T cells, we next used an *in vitro* suppressor assay to measure the function of Tregs in the presence or absence of DT. Deleting PD1^+^ Tregs significantly abrogated the suppressor function of Tregs in *in vitro* experiments (**Fig. 6a-b; Extended Fig. 9f**). Next, we established an experimental murine model of T cell driven colitis using our previously optimised *in vivo* DTR protocol^14^. Here, *Rag1^-/-^* mice were monitored for weight loss in the absence and presence of DT (**Fig. 6c**). Weight loss and decreased colon length was significant in animals treated with DT (**Fig. 6d-**e). On performing immunobiology of the LPLs and the MLNs, we found a significant decrease in FoxP3^+^ Treg frequency as expected in the DT cohort, unexpectedly we also found a significant increase in the exFoxP3 Treg frequency both in the LPLs and the MLNs (**Extended Fig.9g**; **Fig. 6f-h**). On measuring IL10 expression, given its role in colitis, we found a significant decrease in Treg-derived IL10 in the DT-treated cohorts (**Fig. 6i-k**). Next, we explored the phenotype of ex-FoxP3^+^ cells to determine whether it is simply a downregulation of IL10 cytokine or an increase in IL17 and IFNγ which further contributes to colitis. We found a significant increase in IL17 production within the exFoxP3 Tregs, along with a significant increase in IFNγ^+^IL17^+^ phenotype (**Fig. 6l-m**). Next, we measured the absolute cell numbers of IL17^+^and IFNγ^+^IL17^+^ expressing exFoxP3 cells and found a significant increase in both populations within the MLN and the LPLs (**Extended Fig. 9h)**. We further confirmed this phenotype in the B16F10 melanoma model. Here, animals were inoculated with tumors and then on tumor establishment, congenic CD45.1^+^T cells were adoptively transferred along with Tregs from the *Pd1^fl/fl-DTR-Tdtomato^* mice. DT treated cohorts showed a decrease in tumor burden as expected (**Fig. 6n-o**). Immunobiological analysis of TILs showed a decrease in Treg frequency as expected in the DT cohort (**Extended Fig. 9i**; **Fig. 6p left panel**). Consistent with colitis data, an increase in exFoxP3 cells were noted in the DT cohort (**Fig. 6p right panel**). These data suggest that deleting PD1^+^ Tregs which mark TOX^+^Tregs results in destabilizing the Tregs during inflammation and cancer. To further test this hypothesis, *in vitro* experiments were conducted. Here, PD1^+^ Tregs were significantly enriched in HELIOS, NRP1, CD25 and FoxP3 protein (**Extended Fig. 10a-e**). On sorting PD1^-^ Tregs and culturing them with effector cytokines *in vitro* resulted in a majority of these cells losing Foxp3 expression (**Extended Fig. 10f-g**) further providing evidence for extrinsic control of Treg functional plasticity by PD1^+^ Tregs. Next, we measured the cytokine expression in the FoxP3 compartment and found a significant decrease in IL10 within the TIL Tregs in the presence of DT (**Fig. 6q**). Within the T effector compartment, a significant increase in CD4^+^ T effectors were noted in the cohorts that received the DT (**Fig. 6r-s**). No change in CD8^+^ T cell numbers were seen (**Fig. 6t**). Hence, we conclude that PD1^+^ Tregs predominantly express IL10 and are important in preventing autoimmunity and anti-tumor immunity

**Figure 6:**
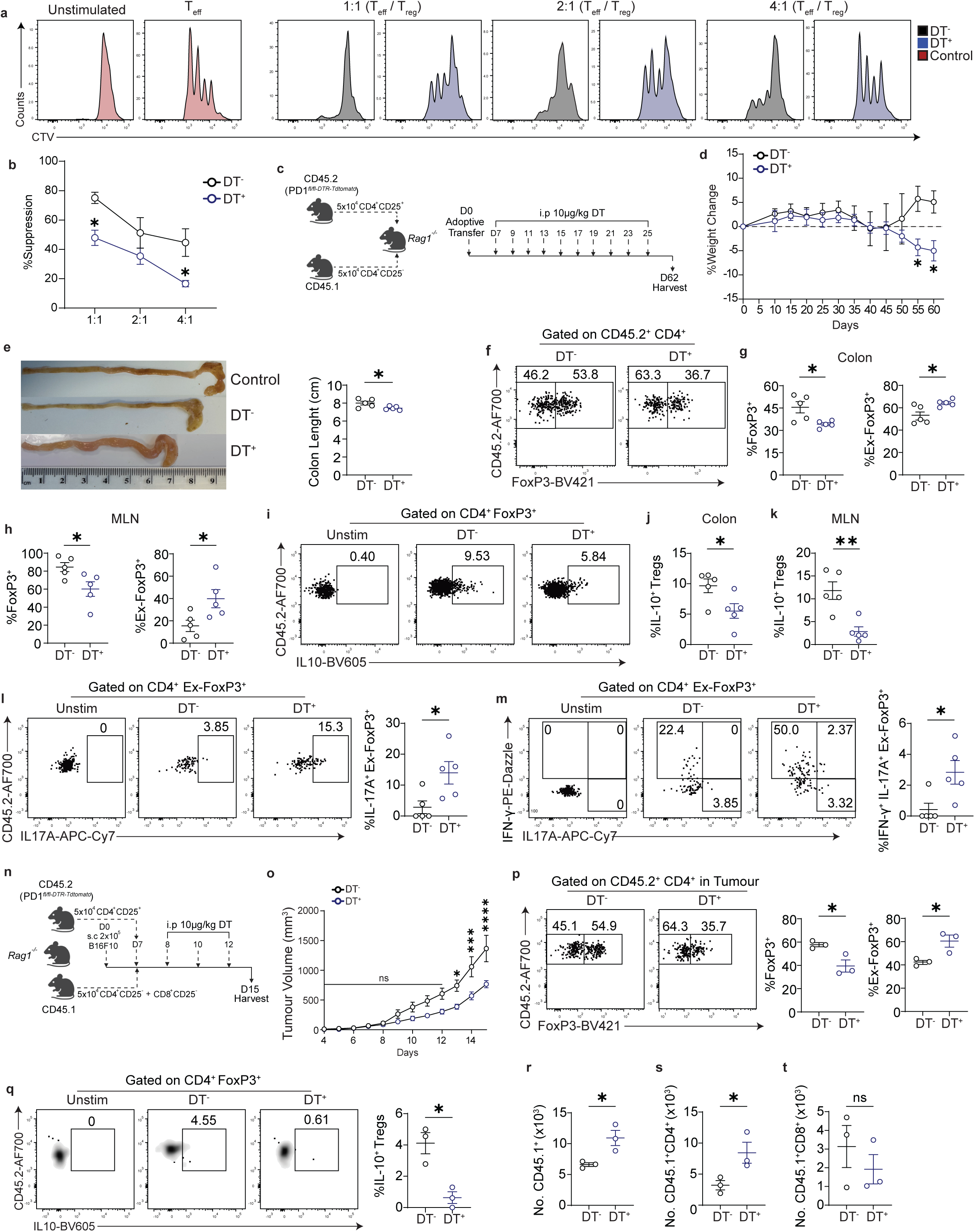
Deleting PD1^+^ Tregs drives experimental autoimmune colitis and enhances anti-tumor immunity. Naïve CD4^+^CD25^-^FoxP3^RFP-^ T cells were cocultured Tregs from B6.*Pdcd1 ^fl/fl-DTR-Tdtomato^Foxp3^cre-ERT2^* mice along with irradiated splenocytes from WT mice. Responder T cells were stained with cell trace violet and Tregs were stained with CFSE. Representative plots of proliferation with and without Tregs; in some cohorts DT was added to the responder T: Treg cultures (**a**). Percent suppression was measured using the division index (**b**). *Rag1^-/-^* mice were reconstituted with naïve congenic CD45.1^+^CD4^+^ T cells plus Tregs from *Pdcd1^fl/fl-DTR-Tdtomato^Foxp3^cre-ERT2^ mice.* Certain cohorts were treated with DT or PBS as shown (**c**). Weight loss was measured (*n*= 5 samples per group) (**d**) and animals were euthanized as per the animal license limit of <15% weight loss from initial weight loss. Length of colon was measured (*n*= 5 samples per group) (**e**). Representative flow plot of Tregs from PBS and DT cohort (**f**), cumulative data showing frequency of FoxP3^+^ and exFoxP3 cells in the LPL (**g**) and in the MLN (**h**). LPLs were stimulated for 4hrs with cytokine stimulation cocktail and then cytokines IL10, IFNγ and IL17 was measured. Representative flow plot of IL-10 in LPL (**i**), cumulative frequency of IL-10 in LPL (**j**), cumulative frequency of IL-10 in MLN (**k**). Representative flow plot of IL17 and cumulative data (**l)**, Representative flow plot of IL17 and IFNγ and cumulative data (**m**). *Rag1^-/-^* mice were reconstituted with B16F10 melanoma. When tumors engrafted at day 7, naïve congenic CD45.1^+^CD4^+^ T cells plus Tregs from *Pdcd1^fl/fl-DTR-Tdtomato^Foxp3^cre-ERT2^ mice.* Certain cohorts were treated with DT or PBS as shown (**n**), tumor volume was measured in both cohorts (*n*=3 samples per group) (**o**), representative flow plot showing frequency of Foxp3 and cumulative FoxP3 frequency and exFoxP3 frequency in both cohorts (**p**). TILs were restimulated with cytokine stimulation cocktail for 4hrs and then IL10 cytokine was measured in both cohorts (**q**). The absolute numbers of CD45.1, (**r**), CD4 (**s**) and CD8 (**t**) in both cohorts within the TILs. Each dot represents a mouse, data shown are Mean±SEM, a un-paired student t test and two-way ANOVA with Sidak’s multiple comparison (**d, o**) was performed. \**P* ≤ 0.05, \*\**P* ≤ 0.01, \*\*\**P* ≤ 0.001, \*\*\*\**P* ≤ 0.0001.

### Blocking PD1 increases TOX^+^ Tregs in human melanoma TILs

TOX expression in human Tregs in cancers has been recently reported^31^ ^30^, but the outcome of anti-PD1 treatment in this cell population is unknown. Using publicly mined dataset, we explored the phenotype of PD1^+^TOX^+^ Tregs in treatment naïve melanoma (**Fig. 7a-b; Supplementary Table 2-4).** PD1^-^ and PD1^+^ Treg subsets were present in this dataset. Furthermore, PD1^+^ subset was enriched in TOX expressing Tregs (**Fig. 7c-d**). On further characterizing this PD1^+^ Treg TILs in human melanoma, we found a significant increase in *IL10* mRNA within this subset and expression of *HAVCR2, MKI67* and *BCL6* although these did not reach significance (**Fig. 7e)**. Next, we analyzed melanoma samples that were treatment naive or treated with anti-PD1 antibody (**Fig. 7f-g**). After anti-PD1 treatment, the total frequency of TOX^+^ Tregs significantly increased, no change in TCF1^+^TOX^+^ Treg subset was noted and a significant increase in TCF1^-^TOX^+^ cell frequency was found (**Fig. 7h**). On further analyzing the TCF1^-^TOX^+^ Treg subset, a significant increase in *HAVCR2* was noted (**Fig. 7i**). Further bulk mRNA analysis of each individual sample showed a significant increase in *IL10* mRNA per individual (**Fig. 7j**). No change in other markers namely *HAVCR2, MKI67* and *BCL6* was noted. These data indicate that anti-PD1 treatment can increase TCF1^-^TOX^+^ Tregs that also express *HAVCR2* and *IL10* which may contribute towards an immunosuppressive microenvironment and anti-PD1 treatment resistance.

**Figure 7:**
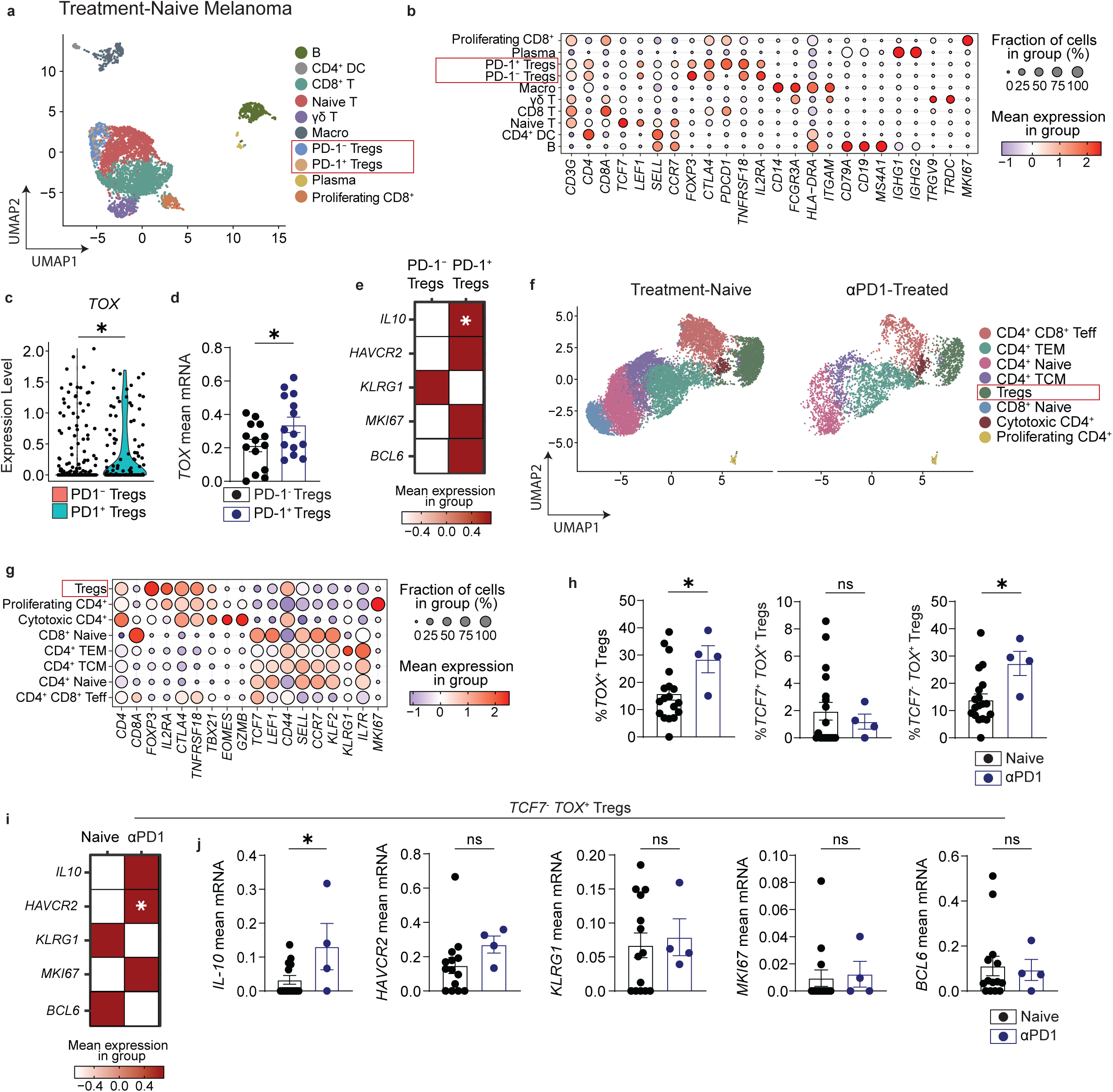
PD1 marks TOX^+^ Tregs in human melanoma and blocking PD1 increases the frequency of TOX^+^ Tregs. Public dataset on treatment naïve melanoma and anti-PD1 treated melanoma samples were mined and then used for analysing the frequency of *PD1^+^TOX^+^*Treg subsets. UMAP of treatment naïve melanoma TILs (**a**), Dot plot highlights markers used to define the various immune cell subsets in the UMAP (**b**), violin plot showing expression of *TOX* within *PD1^-^* and *PD1^+^* Tregs (**c**), mean mRNA expression of *TOX* in individual samples in *PD1^-^* and *PD1^+^*Tregs (**d**). Heatmap showing mean expression of *IL10, HAVCR2, KLRG1, MKI67, BLC6* (**e**). UMAP of melanoma samples consisting of treatment naïve and anti-PD1 treated cohorts (**f**). Dot plot showing markers used to define populations within the dataset (**g)**. Frequency of *TOX^+^* Tregs, *TCF7^-^TOX^+^* Tregs and *TCF7^+^TOX^+^* Tregs (**h**). Heatmap showing mean expression of *IL10, HAVCR2, KLRG1, MKI67* (**i**). Mean mRNA expression of *IL10, HAVCR2, KLRG1, MKI67* within *TCF7^-^TOX^+^*Tregs in individual samples (**j**). Each dot represents an individual sample, data shown are Mean±SEM, a Wilcoxon rank sum test was performed for the mean expression visualised as heatmap, paired (**d**) and un-paired (**j**) student t test was performed for bulk mRNA analysis. \**P* ≤ 0.05, \*\**P* ≤ 0.01, \*\*\**P* ≤ 0.001, \*\*\*\**P* ≤ 0.0001.

## Discussion

The majority of peripheral Tregs constitutively express the inhibitory co-receptor CTLA4, but PD1 is expressed on a specific subset of peripheral Tregs, the significance of this distinct subset is unclear. In this report we elucidate three key aspects of PD1 in Treg biology. (1) we demonstrate that PD1 expressing Tregs are enriched in the transcription factor TOX in steady state, (2) PD1 not just marks TOX^+^ Tregs but intrinsically controls their frequency and absolute numbers in steady state and (3) deleting PD1^+^ Tregs using a previously undefined mouse model of PD1-DTR highlights the function of this subset in protecting against autoimmunity and dampening anti-tumor immunity. To date, the role of PD1^+^ Tregs, expression of TOX in Tregs in steady state and deletion of PD1^+^ Tregs as a therapeutic approach has not been investigated or reported. Our data suggests that PD1^+^ Tregs constitute a distinct Treg subset which play a major role in controlling autoimmunity and anti-tumor immunity. PD1 expression is confined to a specific subset that marked both TOX and TCF1 in the periphery. Blocking PD1 or germline deletion significantly increased the TOX^+^ Tregs. Our finding that TOX is expressed in Tregs has been previously reported within certain cancers ^30^ but in-depth phenotyping of TOX in Tregs in steady state is yet to be reported. Herein lies the importance of this work whereby we investigated if TOX behaves similar to its role in CD8^+^ T cells or CD4^+^ T effectors in Tregs.

TOX has emerged as a key transcription factor in driving a distinct CD8^+^ T cell differentiation programme that leads to Tex cells^23, 24^. In CD4^+^ T cells, TOX has recently been shown to drive Th1 and Tfh1 effector function^29^. Our data highlights a key role for TOX in Treg effector function whereby deletion of this population results in dampened IL10 production, which results in increased autoimmunity and reduced tumor burden. Taken together, we propose that similar to its role in CD4^+^ T effectors, TOX is necessary for the function of a distinct peripheral Treg subset. This population of TOX^+^ Tregs are controlled by PD1 in the periphery.

TOX is induced by T cell receptor activation ^29^ of NFAT2 signaling, can also induce the expression of PD1^36^. Further, TOX itself can control the expression of inhibitory coreceptors including PD1. Hence one can postulate that peripheral Tregs are constantly receiving T cell receptor stimulation, driven by autoantigen as part of their peripheral immune surveillance ^29^. This in turn leads to TOX and PD1 expression on Tregs in steady state unlike CD8^+^ T cells that express TOX under chronic viral ^24^ or tumor microenvironments. Consistent with this, PD1 also marked TCF1^+^TOX^+^ subset in Tregs. TCF1 is expressed on naïve Tregs, which on activation can upregulate TOX, that ultimately results in the loss of TCF1. In contrast with the induction of PD1 by TOX, we found that PD1 inhibited TOX^+^Treg frequency, presumable due to its ability to inhibit T cell receptor signaling, thereby preventing an expansion of TOX^+^ Tregs that can ultimately dampen the function of CD8^+^ and CD4^+^ T effectors in the periphery. Indeed, psuedotime trajectory analysis of PD1^+^ Treg subsets does highlight a trajectory that follows the proposed pathway with the double positive TCF1^+^TOX^+^ Tregs transitioning into TCF1^-^TOX^+^ subsets in tissues.

Using a tamoxifen inducible Cre limited to FoxP3^+^ Treg, we found that deletion of PD1 within the Treg compartment led to enhanced tumor cell growth^14^. Thus, anti-PD1 can result in the expansion of Tregs within the TME thereby enhancing tumor growth and driving checkpoint resistance. By contrast, when we deleted PD1^+^ Treg using a DTR model, tumor growth was inhibited. Both *in vitro* and *in vivo* experiments demonstrate that deleting PD1^+^ Tregs can abrogate Treg cell suppression, enhance tumor immunity and dampen protection against colitis. Deleting Tregs within the TME by targeting the IL2 receptor ^4^ ^37^or chemokines such as CCR8^38^ has been postulated. Indeed, pH driven targeted deletion is also being extensively investigated which ultimately aims to delete Tregs within the TME. The shortcomings of such approaches include deleting all Tregs within the TME. The shortcomings of such approaches include deleting all Tregs within the TME, even those that could be enriched for protective peripheral tissue immunity thereby enhancing anti-PD1 related toxicity ^39^. By specifically deleting PD1^+^ Tregs, one can overcome this hurdle whereby the majority of Treg subsets are protected while the specific effector Treg subsets are deleted.

Anti-PD1 can also reinvigorate CD8^+^Tpex but this effect is short lived within the CD8 compartment due to stable exhausted epigenetic phenotype. One could hypothesize that during anti-PD1 treatment while the short lived CD8^+^ T cell boost occurs without epigenetic correction of CD8^+^ Tex, a concurrent expansion of suppressive TOX^+^ Tregs within the TME also occur since they respond to anti-PD1. If this is the case, the competitive survival of both these populations can determine the ultimate outcome of tumor immunity.

Finally, our data also highlights a function for PD1^+^ Treg population in extrinsically controlling functional plasticity of PD1^-^Tregs. On deletion of PD1^+^ Tregs in tumor models, colitis and *in vitro* assays, we found a significant increase in ex-FoxP3 populations. PD1 signaling has an intrinsic role in maintaining Foxp3 expression in Tregs and Th1 cells through several mechanisms^11, 12, 13, 14, 40, 41^ Perhaps deleting PD1^+^ Treg population, highlights the phenotype of the Tregs that are left behind which then preferentially expand and lose their identity.

In summary, this study describes the immunobiology of PD1^+^ Treg subsets in steady state and in disease conditions. We show PD1 marks a population of TOX-expressing Tregs which contribute towards pro-tumor immunity and prevents autoimmune colitis. Deleting this population with a novel PD1-DTR model restores anti-tumor immunity. This data can contribute towards the tailoring of PD1 based therapeutics both in cancer and autoimmunity.

## Materials and Methods

### Animals

C57BL/6, CD45.1 C57BL/6, *Rag1^-/-^* , BALB/c mice were purchased from Charles River Laboratories while *Foxp3^RFP^* and *Foxp3^ERT^*^2^*^-Cre-eGFP^*mice were purchased from Jackson Laboratory. *Pdcd1^-/-^ (Pd1^-/-^)-*C57BL/6 were provided by Dr Lieping Chen (Yale University School of Medicine, New Haven, CT) and were further bred to *Foxp3^RFP^*mice to generate *Pd1^-/-^Foxp3^RFP^* mice. *Pd1^fl/fl-DTR-Tdtomato^*mice were engineered by Ozgene and were further bred to *Foxp3^Ert2-Cre-eGFP^*to generate *PD1^fl/fl-DTR-Tdtomato^Foxp3^Ert2-Cre-eGFP^*mice. To inducibly ablate PD1 expressing cells, *Pd1^fl/fl-DTR-Tdtomato^*mice were given 10ug/kg of diphtheria toxin from *Corynebacterium diphtheriae* (Sigma-Aldrich) in PBS up to 10 dosages via intraperitoneal injection (IP). *Pd1^fl/fl-DTR-Tdtomato^* mice were further bred to *Foxp3^Ert2-Cre-GFP^* to generate *Foxp3^Ert2-Cre-GFP^Pd1^fl/fl-DTR-Tdtomato^* mice that can selectively eliminate PD1 in FoxP3-expressing regulatory T cells (Tregs). To induce Cre-mediated PD1 excision, *Foxp3^Ert2-Cre-GFP^Pd1^fl/fl-DTR-Tdtomato^* mice were given 1mg tamoxifen (Sigma-Aldrich) in corn oil (Sigma-Aldrich) up to 5 dosages via IP. To inhibit PD1 signalling, BALB/C mice were given 0.2mg anti-PD1 (clone: RPM1:14) or IgG up to 3 dosages via IP. All mice were bred and maintained in a pathogen-free facility at Newcastle University under a home office-approved project license. All experimental animals were 8 to 12 weeks of age. All murine experimental procedures were performed at Newcastle University incorporating the NC3R guidelines for animal research, and the data are presented according to the ARRIVE guidelines.

### Reagents

Mouse complete media was made up of DMEM (Gibco) supplemented with 10% FBS, 2mM L-glutamine (Corning), 0.1mM non-essential amino acids, 1mM sodium pyruvate (Sigma-Aldrich), 50µM 2-mercaptoethanol and penicillin and streptomycin (100 units/ml penicillin; 100µg/ml streptomycin, Corning). Melanoma culture media was made up of DMEM (Gibco) supplemented with 10% FBS and penicillin and streptomycin (100 units/ml penicillin; 100µg/ml streptomycin). Flow cytometry staining (FACS) buffer was made up of PBS supplemented with 0.5% BSA (Sigma-Aldrich) and 0.01% sodium azide. Miltenyi buffer was made up of PBS supplemented with 0.5% BSA and 2mM EDTA (Sigma-Aldrich).

### Flow Cytometry

Single-cell suspensions from indicated organs were stained with Live/Dead fixable dead cell stain kit as per manufacturer’s instruction (Invitrogen). Surface staining was performed for 30 min at 4°C in FACS buffer with the following antibodies: CD4 (GK1.5, Biolegend), CD44 (IM7, Biolegend), CD45.1 (A20, Biolegend), CD45.2 (104, Biolegend), CD62L (MEL-14, Biolegend), CD8α (53-6.7, Biolegend), Nrp1 (3E12, Biolegend), PD1 (29F.1A12, Biolegend), CD73 (TY-11.8. Biolegend), CD127 (A7R34, Biolegend), TIM-3 (RMT3-23, Biolegend), CD25 (PC61, Biolegend) CD16/32 (93, Biolegend). For intracellular cytokine staining, cells were stimulated with cell stimulation cocktail containing protein transport inhibitors (eBioscience) for 4 h at 37°C. Cells were then fixed and permeabilized using FoxP3 Transcription Factor Staining Buffer Set (eBioscience) as manufacturer’s instruction. Intracellular staining was performed overnight at 4°C in 1x permeabilization buffer (eBioscience) with the following antibodies: BCL6 (7D1, Biolegend), TOX (TXRX10, Invitrogen), TCF1 (S33-966, BD), FoxP3 (FJK-16S, eBioscience), Helios (22F6, Biolegend), IFNγ (XMG1.2, Biolegend), IL10 (JES5-16ES, Biolegend), IL17A (TC11-18H10.1, Biolegend) and CD16/32 (93, Biolegend).

Flow cytometry was performed using a BD LSR Fortessa X20. Sort purification was performed on a BD FACSAria Fusion. All data were analyzed using FlowJo 10 (Tree Star). Graphs and statistical analyses were done with Prism 10 (GraphPad Software).

### Diphtheria toxin dose optimisation

Splenocytes were obtained from *Foxp3^Ert2-Cre-GFP^Pd1^fl/fl-DTR-Tdtomato^*mice. 2×10^5^ splenocytes were stimulated with soluble anti-CD3 (5 μg/mL, Clone: 145-2C11). Cultures were treated with diphtheria toxin at indicated concentrations. Cells were cultured in complete media for 24, 48 and 72 h at 37°C with 5% CO_2_ and PD1 expression was assessed on flow cytometry.

### Treg suppressor assays

Treg suppressor assay was performed as previously published ^42^. Naïve T cells were obtained from *FoxP3^RFP^* mice and Tregs from *Foxp3^Ert2-Cre-GFP^Pd1^fl/fl-DTR-Tdtomato^*mice. Sorted CD4^+^ T naïve cells were stained in PBS with cell trace violet (CTV) and Tregs were stained with CFSE. 5×10^4^ Teff and the indicated ratios of sorted Tregs from *Pd1^fl/fl-DTR-Tdtomato^*mice were stimulated using 0.5μg/ml anti-CD3 (Clone: 145-2C11, Biolegend) along with irradiated splenocytes from C57BL/6 WT mice as antigen-presenting cells (APCs) using a 4:1 ratio of APCs to Teff cells. Cultures were treated with diphtheria toxin or PBS (0.5ug/ml). Cells were cultured in complete media for 72 h at 37°C with 5% CO_2_ and proliferation was assessed on flow cytometry.

### Treg Plasticity Experiments

CD90.2^+^ T cells were enriched from *FoxP3^RFP^* murine splenocytes using CD90.2 Microbeads (Miltenyi Biotec according to the manufacturer’s instructions). Enriched cells were stained with CD4-AF700 (Clone: GK1.5), and anti-CD16/32 (Clone: 93) for 15 min at 4°C. DAPI was added immediately prior to sorting, and live and PD1^tdtomato-^CD4^+^FoxP3^GFP+^ Treg cells were sorted on a BD FACSAria^TM^ Fusion cell sorter. Sorted Tregs were plated at 6×10 cells/mL in mouse complete media with plate-bound anti-CD3 (5 μg/mL, Clone: 145-2C11), soluble anti-CD28 (2 μg/mL, Clone: 37.51) and cultured for 6 days under one of the following polarising conditions (all cytokines from Miltenyi Biotec): IL-4 (5 ng/mL), IL-6 (20 ng/mL), IL-12 (20 ng/mL and IL-18 (20 ng/mL) or IL-1β (50 ng/mL) and IL-23 (50 ng/mL). IL-2 was supplemented on days 3 and 5, and IL-4 was re-added on day 3 and 5. Cells were harvested on day 6 and FoxP3 expression assessed by flow cytometry as a measure of Treg lineage stability.

### Tumor model

B16 melanoma and MC38 was set up using established protocols^11, 43^. Tumor cells were cultured in complete mouse tumor media at 37 °C with 5% CO_2_. Mice were inoculated with 2×10^5^ B16F10 melanoma cells via subcutaneous injection into the flank as previously reported ^44^. Tumor volume was measured daily from day 5 post inoculation and calculated by the following equation: tumor volume = π/6 x 0.5 x length x width^2^ as described previously ^45^For Treg adoptive transfer experiment, *Rag1^-/-^* mice were re-constituted with 5×10^4^ sorted CD4^+^CD25^-^ and CD8^+^CD25^-^ from CD45.1 WT mouse as Teff cells along with 5×10^4^ CD4^+^CD25^+^ as Tregs from *Pd1^fl/fl^Foxp3^ERT2cre^*mice via intravenous injection at day 7 post inoculation. Cohorts either received PBS or diphtheria toxin (DT) every other day, with a total of 3 doses (10ug/kg). Mice were euthanized when tumor volume reached 800mm^3^-1500mm^3^.

### Experimental Autoimmune Colitis

*B6.Rag1^-/-^* mice were reconstituted with B6. CD45.1^+^CD4^+^CD25^-^ (T effectors) populations (5×10^4^) as previously described^46^ along with flow sorted B6. CD4^+^CD25^+^ cells from *Pd1^fl/fl^Foxp3^ERT2cre^*mice. Mice were treated with either diphtheria toxin or PBS every other day in a total of 10 doses (10ug/kg). Mice were weighed weekly and loss of body weight was used as an indicator of colitis Mice were euthanized when weight loss reached <15% in accordance with Home Office allowed clinical endpoint. Immune endpoints were measured in the MLN and in LPL. LPL were isolated as previously described^46^, both lymphocytes from MLN and LPL were stimulated with cytokine stimulation cocktail for 4hrs and then effector cytokines were measured by intracellular flow cytometry.

### Dextran sulfate sodium (DSS)-induced colitis

Colitis was induced in *Foxp3^RFP^* mice by oral administration of 3% (w/v) Dextran Sulfate Sodium Salt (DSS) (Thermo Fisher Scientific) in drinking water for 6 days. Weight loss, stool scoring and presence of blood was recorded daily. Following treatment, mice were sacrificed once the blood was present in the stool, and colons were harvested. Colons were measured and then washed with colon wash buffer (PBS supplemented with 20mM HEPES (Merck), 2% FBS (Biosera) and 2-mercaptoethanol (Sigma Aldrich), then digested with 0.5mg/ml DNase (Merck) and 0.25mg/ml Liberase (Merck) for 15min at 37°C. Lymphocytes were separated using a Percoll gradient (Merck), washed with complete mouse media and processed for flow cytometry.

### ScRNA-seq dimensionality reduction

Publicly available mouse Treg scRNA-seq dataset was pre-processed, and the dataset was retrieved from the National Centre for Biotechnology Information (NCBI) Gene Expression Omnibus (GEO) database. In RStudio (R v.4.5), using Seurat package (v.5.5.1), for each mouse, cell barcodes were matched with corresponding features, and tissue origin was assigned based on antibody hashtag signals when the hashtag count was 30 times higher than the others as described in the original study^35^. Cells were filtered to include those with >200 detected genes and <5% mitochondrial RNA. Expression matrices were log normalized and top 2,000 variable features were identified. Principal component analysis (PCA) was performed for dimensionality reduction (npcs= 50). Harmony (v.2.0.5) was applied on PCA embeddings using each mouse-tissue to remove the batch effect. The top 20 principal components (PCs) were used to compute UMAP. FindNeighbors function (dims= 1:20, reduction= “harmony”) was used to create k-nearest neighbor graph (KNN). Louvian clustering (resolution= 0.4) was used. Two clusters with multiple lineage signals were identified and were removed. Remaining cells were re-clustered using 10 Harmony-corrected dimensions and Louvain clustering at resolution 0.4.

For mining of publicly available human melanoma scRNA-seq datasets^47, 48, 49^ were pre-processed and processed gene expression data were obtained from the GEO and Zelondo databases. In Naïve human melanoma dataset, samples subjected to therapy (Post) were removed, remaining samples were log-normalized, and 2,000 most variable genes were identified. In naïve-anti-PD1 human melanoma dataset, samples which received combination therapies were removed, and 2,000 most variable genes were identified. In both datasets, PCA analysis was performed for dimensionality reduction (npcs= 50). For Naïve human melanoma dataset, 15 PCs were used to create UMAP embedding, followed with KNN graph generation and Louvian clustering at resolution 0.2. In naïve-anti-PD1 human melanoma dataset, Harmony was applied using samples (Patient_ID), top 10 PCs were used to compute UMAP and FindNeighbors function (dims= 1:10, reduction= “harmony”) was used to generate KNN graph. Louvian clustering (resolution= 0.4) was used.

### ScRNA-seq Analysis

In mouse Treg scRNA-seq analysis, 7 Treg clusters were identified (Naïve, *Nrp1^+^, Pdcd1^+^ Tcf7^+^ Tox^+^, Pdcd1^+^ Tcf7^−^ Tox^+^,* Tr17, ISG^+^, Proliferating). In Naïve human melanoma TIL scRNA-seq analysis, 9 clusters were identified (B, CD4^+^ DC, CD8^+^ T, Naïve T, γδ T, Macro, Tregs, Plasma and Proliferating CD8^+^). In naïve-anti-PD1 human melanoma TIL scRNA-seq analysis, 8 clusters were identified (CD4^+^ CD8^+^ Teff, CD4^+^ TEM, CD4^+^ TCM, Tregs, CD8^+^ Naïve, Cytotoxic CD4^+^, Proliferating CD4^+^). Clusters were annotated using defined marker gene expression for each cluster that was obtained using FindAllMarkers function (method= “Wilcoxon” with Bonferroni correction). scRNAtoolVis package (v0.1.0) was used for JJDotPlot function to visualize gene expression of each cluster. In Naive human melanoma scRNA-seq analysis, previously annotated Treg cluster then was stratified into PD1^+^ and PD1^-^ Tregs by detection of *PDCD1* transcript in the raw RNA count matrix. Tregs with at least one *PDCD1* count were classified as PD1^+^ Tregs and those with zero *PDCD1* counts were classified as PD1^-^ Tregs. In naïve-anti-PD1 human melanoma scRNA-seq analysis, previously annotated Treg cluster then was stratified into *TOX^+^*, *TCF7^+^ TOX^+^, TCF7^-^ TOX^+^*by detection of *TOX* and *TCF7* transcript in the raw RNA count matrix (count>0).

### Gene expression profiling and Pseudotime trajectory analysis

Single cell mRNA expression was visualized using VlnPlot function in the Seurat package. The clusWilcox.test function (method= “ds”) in clusrank (v.1.0.4) package was performed for clustered Wilcoxon rank-sum test. Average mRNA expression of genes of interest within Tregs was summarised using mean function in dplyr (v.1.2.1) package. A two-tailed unpaired Student’s t-test was performed.

Pseudotime trajectory analysis was performed using Slingshot (v.2.16.0) package. Existing cluster annotations and the same first 10 Harmony-corrected dimensions used for clustering. Cluster which was enriched for naïve marker genes (*Tcf7, Bach2, Lef1, Ccr7, and Sell)* was selected as the root for the pseudotime analysis.

## Data availability

Mouse scRNA-seq data used in this paper can be accessed using GEO under accession code GSE266111. Naïve human melanoma and anti-PD-1 treated patients scRNA-seq data can be accessed via GEO under accession code <u>GSE120575</u> and at Zenodo (10.5281/zenodo.18833855) for the Sade-Feldman and Lucca datasets, respectively.

## Code availability

The code used in analysing the datasets can be accessed through GitHub at AkchengelA03/PubliclyMinedscRNAseq.

## Statistical Analysis

All data shown were expressed as individual data points with the line at the mean unless otherwise stated. Error bars on graphs showed the standard error of the mean. Two groups were compared with paired or unpaired Student’s t-test or Wilcoxon signed rank test. Multiple groups were compared using one-or two-way ANOVA analysis corrected by Tukey’s test.

## Author Contributions

A.A conceptualized the idea, performed experiments, analyzed data, wrote, and edited the manuscript; J.X.L performed experiments, provided input into scRNA seq analysis, edited manuscript, K.S performed experiments, C.A, A.L, wrote and edited the manuscript. S.A conceptualized the idea, wrote the manuscript, supervised the study.

## Acknowledgements

The work was funded by an MRC project grant, Leo Foundation Future Leader Award, Lister Prize, the NIHR Newcastle Biomedical Research Centre awarded to the Newcastle upon Tyne Hospitals NHS Foundation Trust, Newcastle University and Cumbria, Northumberland, Tyne and Wear Foundation Trust and LeoFoundation Serendipity Grant. A.A is supported by a British Skin Foundation PhD studentship.

## Extended Figure Legends

**Extended Figure 1: PD1 marks TOX^+^ T regulatory cells in wildtype (WT) mice**

C57BL6 *Foxp3*^RFP^ mice splenocytes were characterised using flow cytometry for PD1^+^TOX^+^ Treg subsets. Representative flow cytometry plot and cumulative data showing corelation between PD1 and TOX in spleen, MLN, liver, lung, small intestine LPL and colonic LPL (**a**). Splenocytes were gated on PD1^negative^ subset, PD1^medium^ and PD1^high^ subsets. Representative plot showing the expression of TCF1 and TOX with the PD1 subset (**b**). Cumulative data of the frequency of TCF1^+^TOX^+^ and TCF1^-^TOX^+^ subsets within the various PD1 Treg subsets in the spleen (**c**), MLN (**d**), liver (**e**), lung (**f**), small intestine (**g**), colon (**h**). Each dot represents a mouse, data shown are Mean±SEM, a paired student t test (**a**) and matched one-way ANOVA with Tukey’s multiple comparison (**c-h**) was performed. \**P* ≤ 0.05, \*\**P* ≤ 0.01, \*\*\**P* ≤ 0.001, \*\*\*\**P* ≤ 0.0001.

**Extended Figure 2: Characterisation of TOX+ T regulatory subsets in mesenteric lymph node and tissues in wildtype mice**

C57BL6 *Foxp3*^RFP^ mice mesenteric lymph nodes and tissues were characterised using flow cytometry of TCF1^+^TOX^+^ and TCF1^-^TOX^+^ T regulatory subsets. Frequency of BCL6, CD73, KLRG1, CD44^+^CD62L^+^ and CD44^+^CD62L^-^ expression in MLN (**a**), frequency of BCL6, CD73, KLRG1, CD44^+^CD62L^+^ and CD44^+^CD62L^-^ expression in liver (**b**), frequency of BCL6, CD73, KLRG1, CD44^+^CD62L^+^ and CD44^+^CD62L^-^ expression in lung (**c**), frequency of BCL6, CD73, KLRG1, CD44^+^CD62L^+^ and CD44^+^CD62L^-^ expression in small intestine LPL (**d**) and the frequency of BCL6, CD73, KLRG1, CD44^+^CD62L^+^ and CD44^+^CD62L^-^ expression in colonic LPL (**e**). Each dot represents a mouse, data shown are Mean±SEM, a paired student t test was performed. \**P* ≤ 0.05, \*\**P* ≤ 0.01, \*\*\**P* ≤ 0.001, \*\*\*\**P* ≤ 0.0001.

**Extended Figure 3: Public mined dataset to characterise TOX+ Tregs in blood, kidney and pancreas**

Publicly available single cell RNA-seq dataset from FoxP3 Thy1.1 mice were reanalysed for the presence of TOX^+^ Tregs in blood, liver, SI LPL, kidney and pancreas. Combined UMAP of Treg subsets in all tissue (**a**), feature plot showing Treg specific markers in all tissue (**b**), Dot plot showing specific markers used to define Treg subsets (**c**), UMAP of specific tissue (**d**), frequency plot showing Tregs subsets in various tissue (**e**). Psuedotime trajectory analysis to identify the lineage trajectory of *TCF7^+^TOX^+^*Treg subsets in pancreas (**f**), LPL (**g**) and kidneys (**h**). Data shown on n=4 mice per organ.

**Extended Figure 4: Pseudotime trajectory analysis of Tregs in pancreas, SI and kidneys**

Publicly available single cell RNA-seq dataset from FoxP3 Thy1.1 mice were reanalysed to determine the gene expression profiles of all the lineages derived from pseudotime analysis. Gene expression profile in all lineages in pancreas (**a**), small intestine LPL (**b**) and kidneys (**c**) across pseudotime are shown. Data shown on n=4 mice per organ.

**Extended Figure 5: Germline deficiency of PD1 increased TOX^+^ Tregs in MLN and tissues**

C57BL6*Foxp3*^RFP^ WT and *Pdcd1^-/-^Foxp3*^RFP^ mice mesenteric lymph nodes and tissues were characterised using flow cytometry of TOX^+^ Treg subsets. Representative flow cytometry plot and cumulative data of TCF1 and TOX expression in Tregs in MLN (**a**), liver (**b**), lung (**c**), small intestine LPL (**d**) and colonic LPL (**e**). Each dot represents a mouse, data shown are Mean±SEM, an un-paired student t test was performed. \**P* ≤ 0.05, \*\**P* ≤ 0.01, \*\*\**P* ≤ 0.001, \*\*\*\**P* ≤ 0.0001.

**Extended Figure 6: Peripheral deficiency of PD1 increased TOX^+^ Tregs in MLN and liver**

B6. *Pdcd1 ^fl/fl-DTR-Tdtomato^Foxp3^cre-ERT2^*mice were treated with either PBS or tamoxifen for 5 days and then mesenteric lymph nodes and tissue were harvested for flow cytometry characterisation. Representative flow cytometry plot and cumulative data of TCF1 and TOX expression in Tregs in MLN (**a**) and liver (**b**). Each dot represents a mouse, data shown are Mean±SEM, an un-paired student t test was performed. \**P* ≤ 0.05, \*\**P* ≤ 0.01, \*\*\**P* ≤ 0.001, \*\*\*\**P* ≤ 0.0001.

**Extended Figure 7: Blocking PD1 using anti-PD1 antibody increased TOX^+^ Tregs**

BALB/c mice were injected with either isotype control or anti-PD1 antibody for 5 days and then splenocytes were harvested for flow cytometry. Representative flow plot of TCF1 and TOX expression within Tregs in the spleen (**a**), cumulative frequency and absolute counts of TCF1^+^TOX^+^ Tregs (**b**), and TCF1^-^TOX^+^ Tregs (**c**). Cells were further characterized for the expression of CD73, KLRG1, CD44^+^CD62L^+^ and CD44^+^CD62L^-^. Representative flow cytometry plots and cumulative data showing the expression of markers in TCF1^+^TOX^+^ Tregs in the isotype control and anti-PD1 treatment (**d-e**). Representative flow cytometry plots and cumulative data showing the expression of markers in TCF1^-^TOX^+^ Tregs in the isotype control and anti-PD1 treatment (**f-g**). Each dot represents a mouse, data shown are Mean±SEM, an un-paired student t test was performed. \**P* ≤ 0.05, \*\**P* ≤ 0.01, \*\*\**P* ≤ 0.001, \*\*\*\**P* ≤ 0.0001.

**Extended Figure 8: TOX^+^ Tregs are marked by PD1 within the B16F10 and MC38 TME**

B6.*Pdcd1^fl/fl-DTR-Tdtomato^Foxp3^cre-ERT2^* mice were inoculated with 2×10^5^ B16F10 tumor cells subcutaneously and tumor growth was monitored **(a)**. Tumors were harvested and flow cytometry characterisation of tumor derived lymphocytes (TILs), tumor draining lymph nodes (tdLN) and splenocytes was performed. Representative flow plots and cumulative frequency of PD1 expression in Tregs within the TILs, tdLN and splenocytes (**b-c**). Frequency of PD1^+^ Tregs in TILs, tdLN and spleen (**d**). B6.*Pdcd ^fl/fl-DTR-Tdtomato^Foxp3^cre-ERT2^*mice were inoculated with 2×10^5^ MC38 tumor cells subcutaneously. Representative flow plots and cumulative frequency of PD1 expression in Tregs within the TILs and splenocytes (**e**) Frequency of PD1^+^ Tregs in TILs and spleen (**f**). Each dot represents a mouse, data shown are Mean±SEM, a paired student t test was performed. \**P* ≤ 0.05, \*\**P* ≤ 0.01, \*\*\**P* ≤ 0.001, \*\*\*\**P* ≤ 0.0001.

**Extended Figure 9: Deleting PD1^+^ Tregs drives experimental autoimmune colitis and enhances anti-tumor immunity**

The deletion of PD1 expressing cells using diphtheria toxin was optimised in *in vitro* cultures. Splenocytes from B6.*Pdcd1 ^fl/fl-DTR-Tdtomato^* mice were cultured *in vitro* with DT (**a**) and then deletion of cells expressing PD1 within CD4^+^ and CD8^+^ was monitored by flow cytometry across time points (**b-e**). Representative flow plots of CFSE stained Tregs in suppressor assay (**f**). *Rag1^-/-^* mice were reconstituted with naïve congenic CD45.1^+^CD4^+^ T cells plus Tregs from *Pdcd1^fl/fl-DTR-Tdtomato^Foxp3^cre-ERT2^ mice.* Certain cohorts were treated with DT or PBS. Gating strategy used to identify congenic T cells and Tregs (**g**). Cumulative data of absolute numbers of IL-17^+^ and IL17^+^IFNγ^+^ ex-FoxP3 cells in MLN and colonic LPL (**h**). *Rag1^-/-^* mice were reconstituted with B16F10 melanoma. When tumors engrafted at day 7, naïve congenic CD45.1^+^CD4^+^ T cells plus Tregs from *Pdcd1^fl/fl-DTR-Tdtomato^Foxp3^cre-ERT2^ mice.* Certain cohorts were treated with DT or PBS. Gating strategy used to identify congenic T cells and Tregs (**i**). Each dot represents a mouse, data shown are Mean±SEM, a matched one-way ANOVA with Sidak’s multiple comparison (**c,e**) and un-paired student t test (**h**) was performed. \**P* ≤ 0.05, \*\**P* ≤ 0.01, \*\*\**P* ≤ 0.001, \*\*\*\**P* ≤ 0.0001.

**Extended Figure 10: Deleting PD1^+^ Tregs drives ex-FoxP3 phenotype in Tregs**

Splenocytes from B6.*Pdcd1 ^fl/fl-DTR-Tdtomato^* mice were flow characterised for the expression of Helios, NRP1, CD25 and Foxp3 in PD1^+^ and PD1^-^ Tregs. Representative flow plot showing expression and cumulative data is shown (**a-e**). Splenocytes were flow sorted for PD1^-^ Tregs and cultured in the presence of anti-CD3, anti-CD28 along with cytokines as indicated. FoxP3 expression was measured at day 6 post culture (**f-g**). Each dot represents an individual experiment, data shown are Mean±SEM, a paired student t test was performed. \**P* ≤ 0.05, \*\**P* ≤ 0.01, \*\*\**P* ≤ 0.001, \*\*\*\**P* ≤ 0.0001.

