## Supplementary figures and images for "Programmed Cell Death-1 marks TOX^+^ T Regulatory Cells that Inhibit Autoimmunity and Anti-Tumor Immunity"

### Figure 1

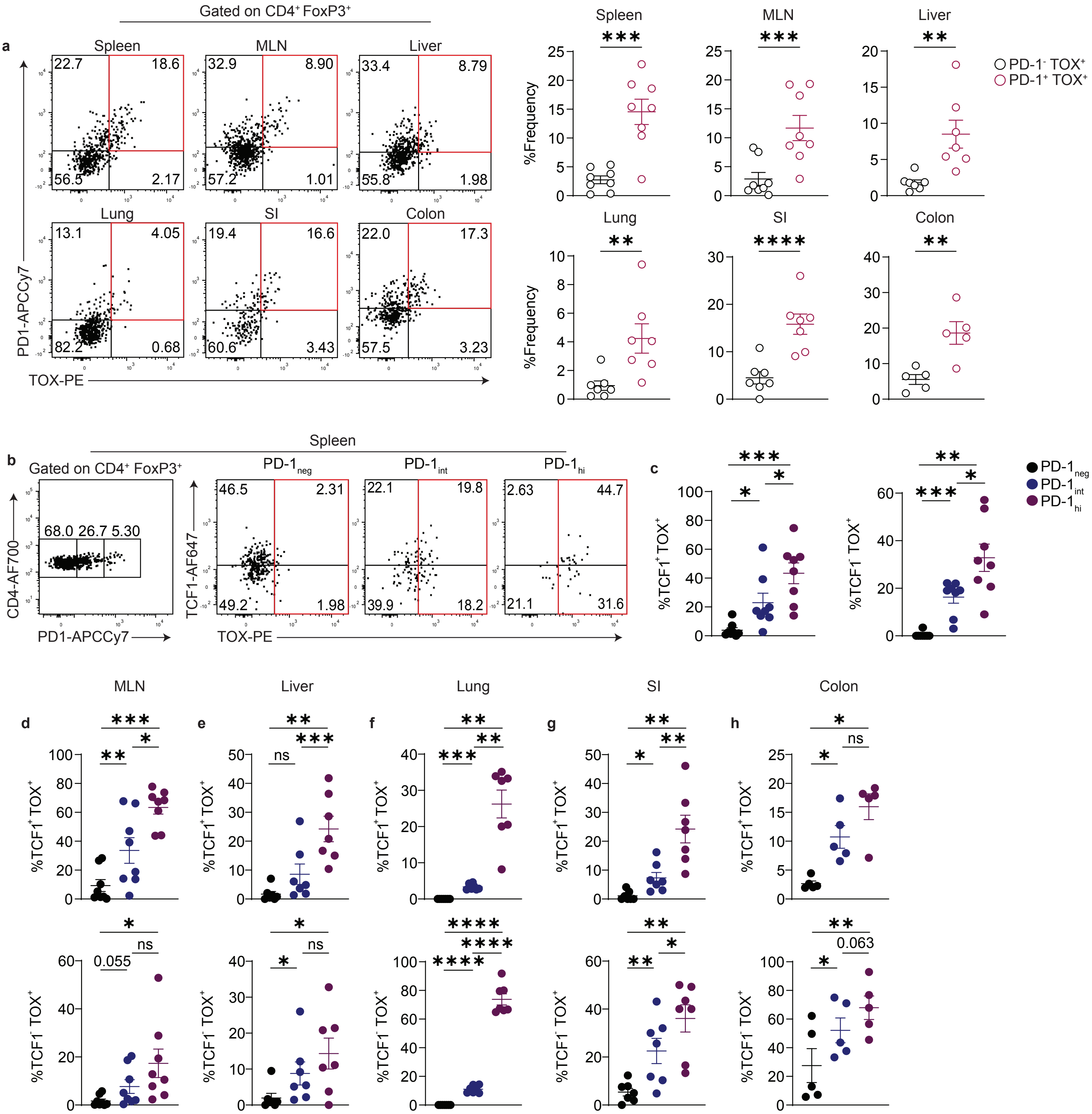


Extended Figure 1

### Figure 1

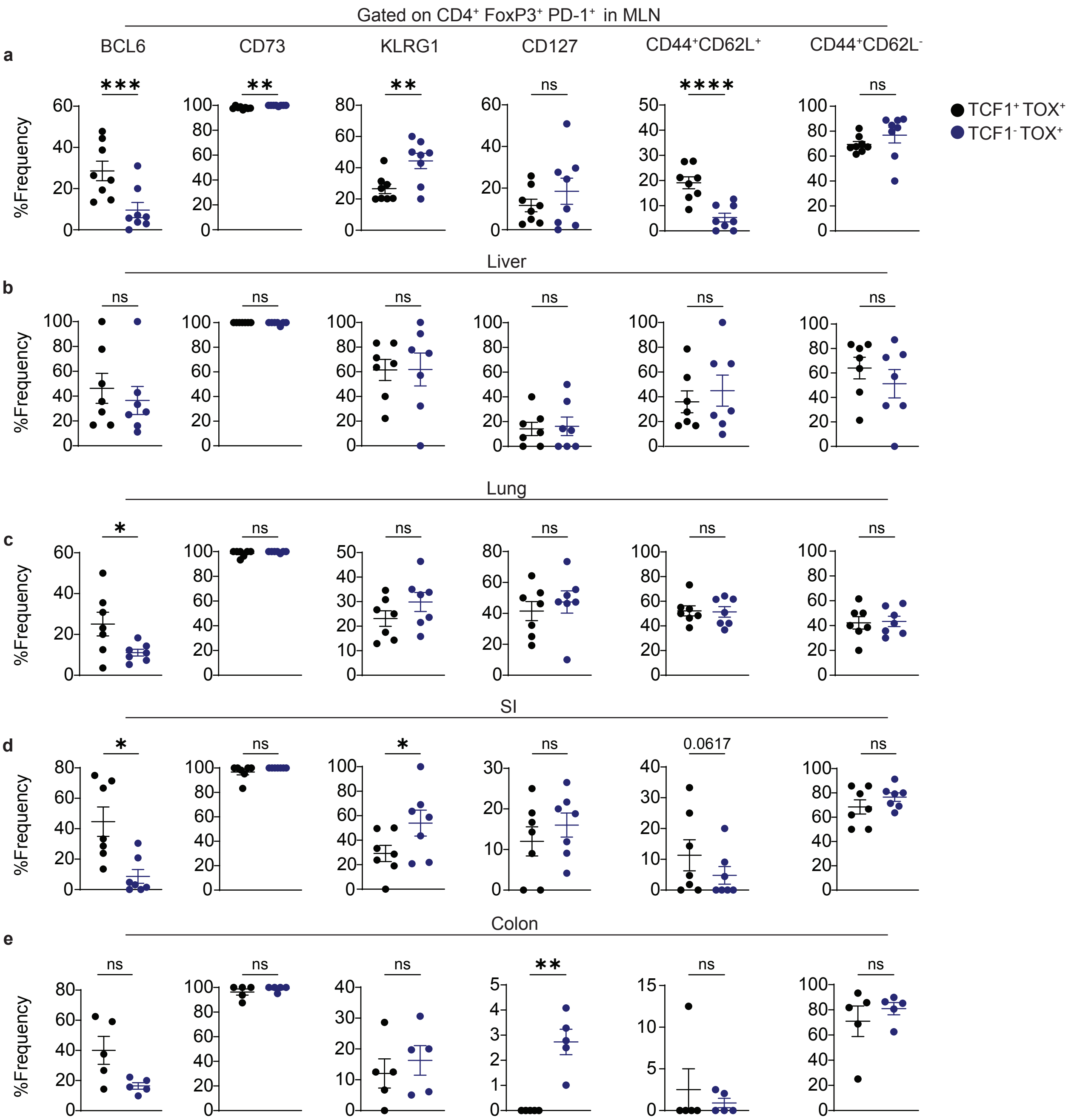


**Extended Figure 2**

### Figure 1

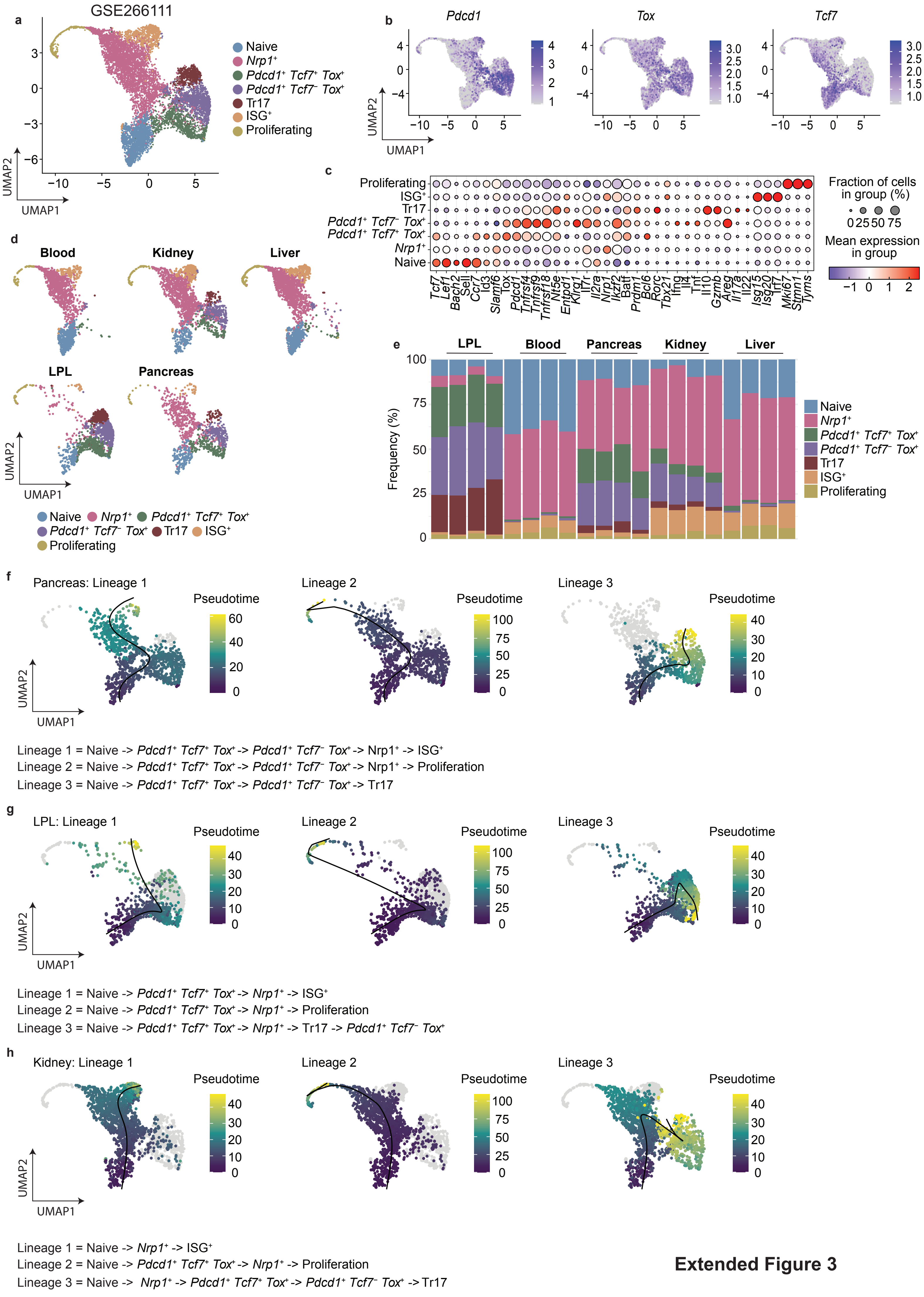


**Extended Figure 3**

### Figure 1

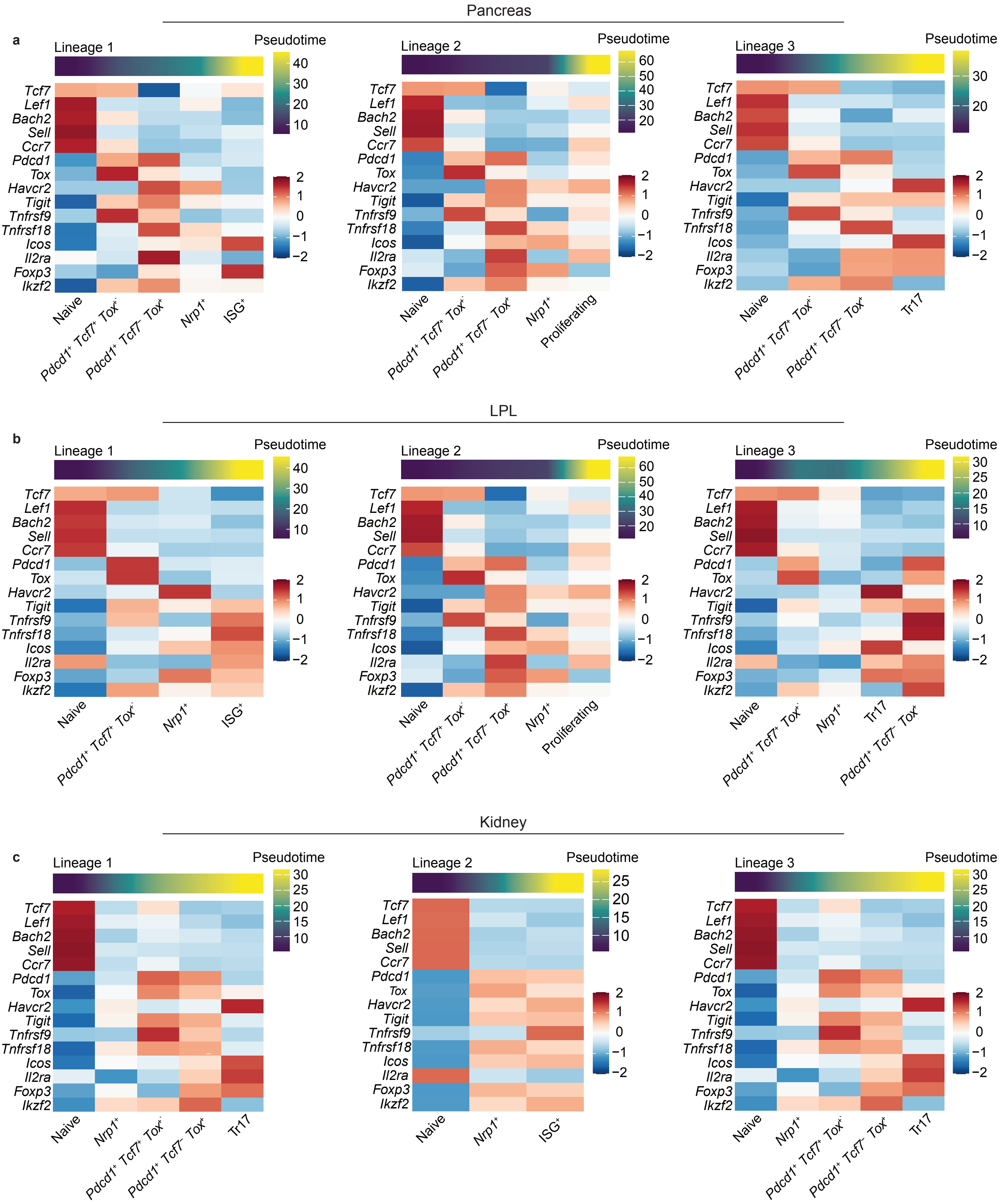


**Extended Figure 4**

### Figure 1

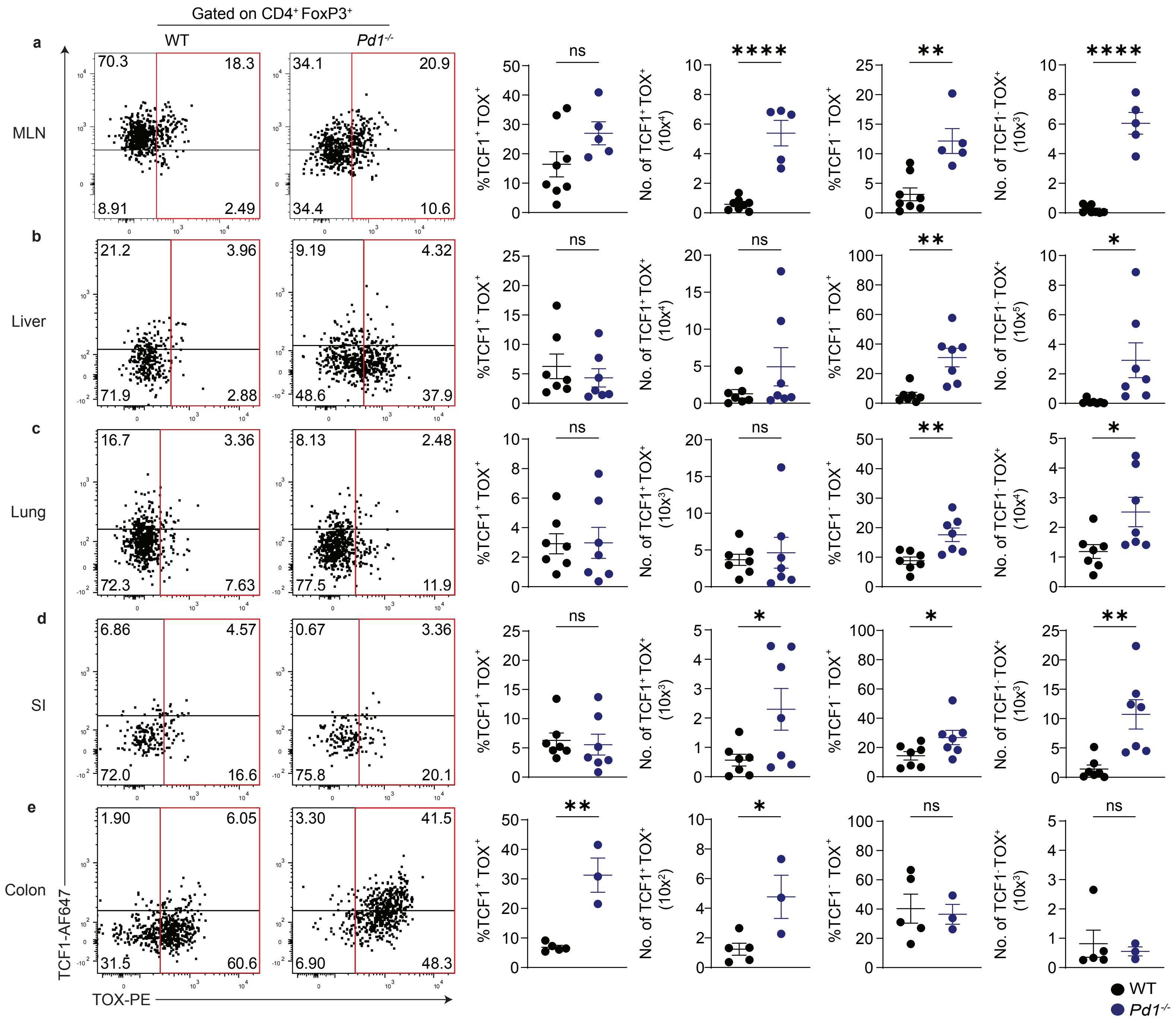


**Extended Figure 5**

### Figure 2

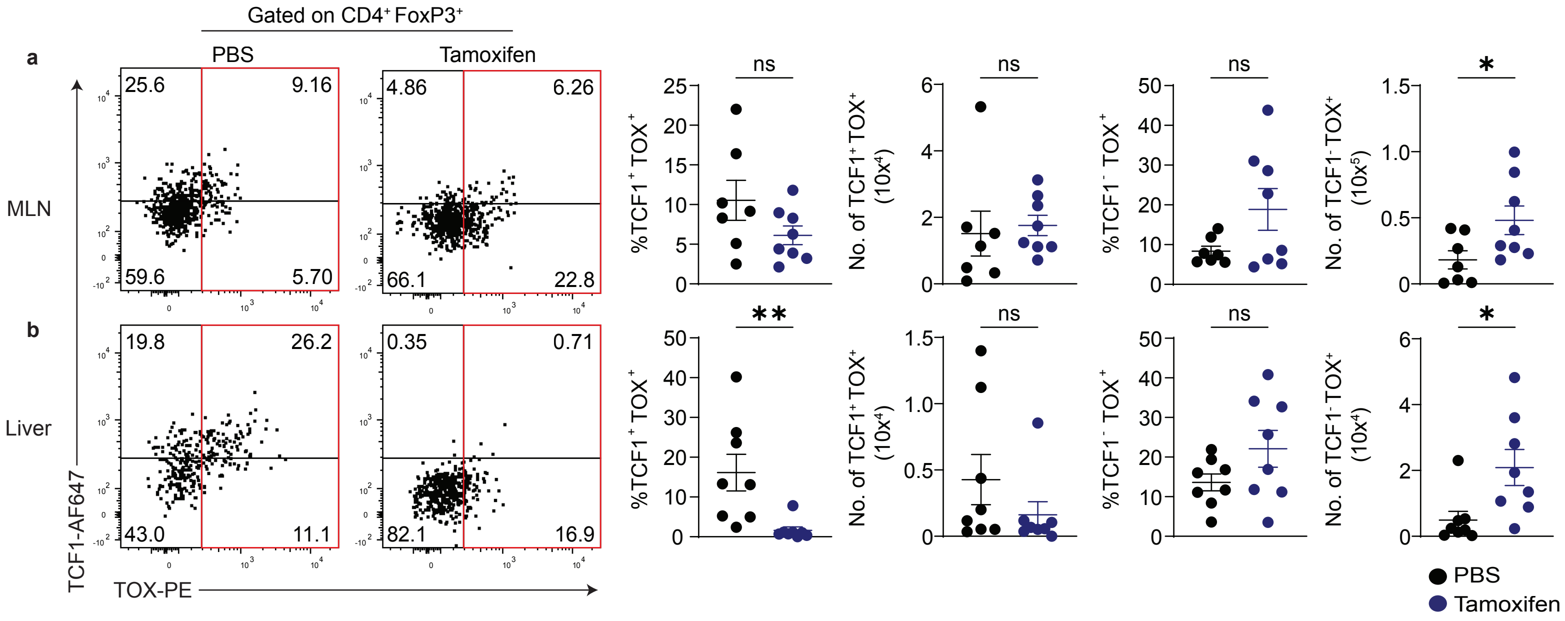


**Extended Figure 6**

### Figure 3

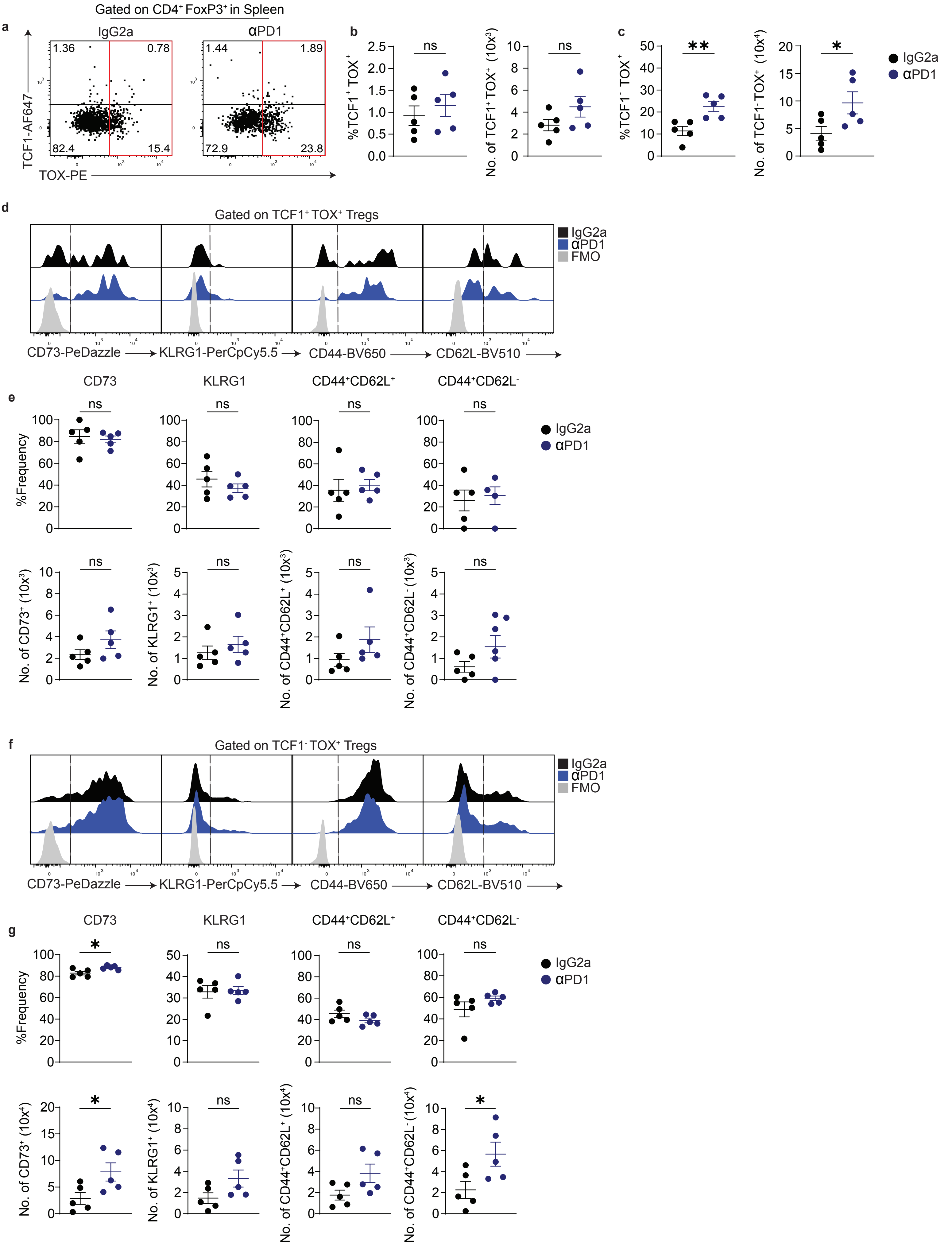


**Extended Figure 7**

### Figure 4

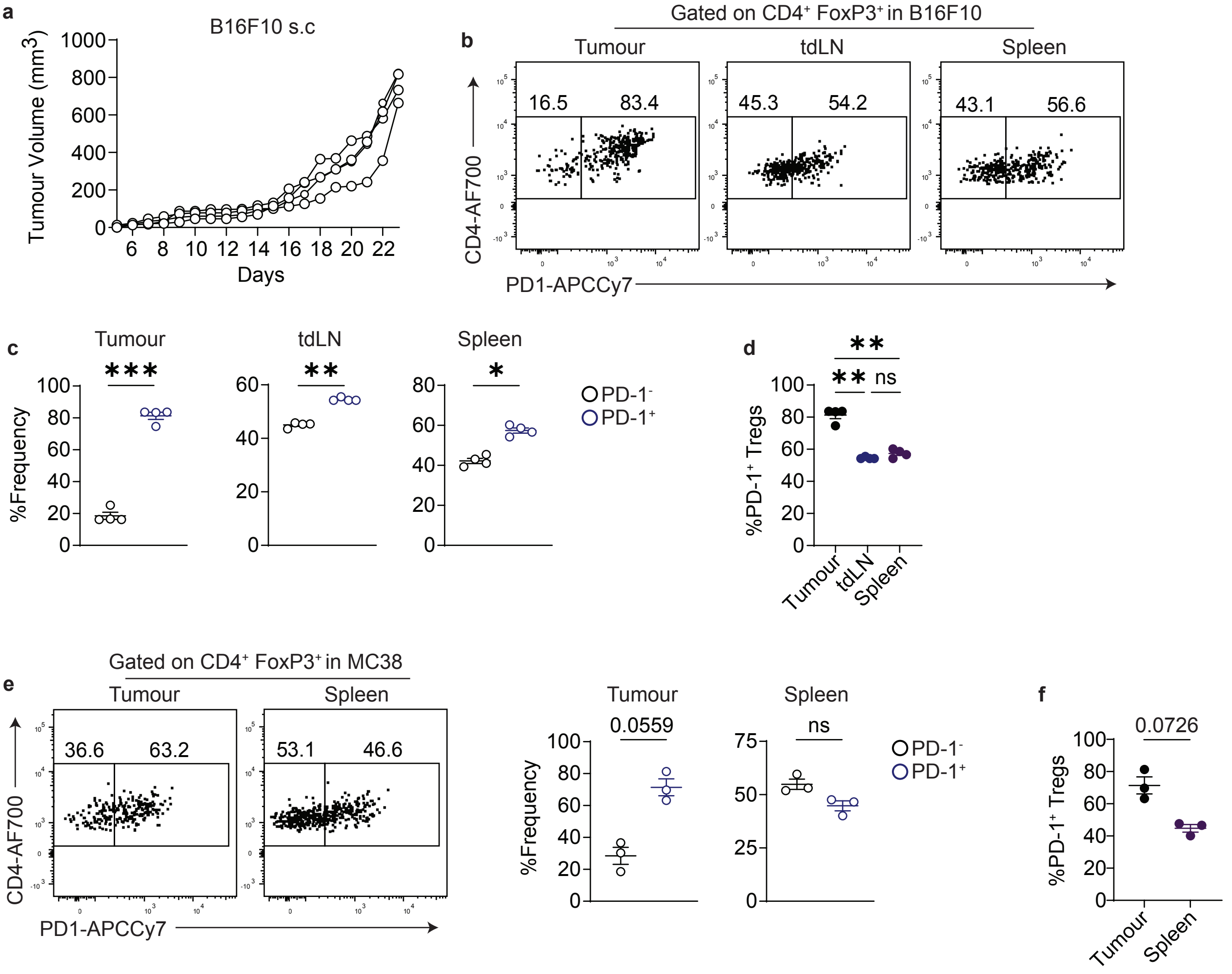


**Extended Figure 8**

### Figure 5

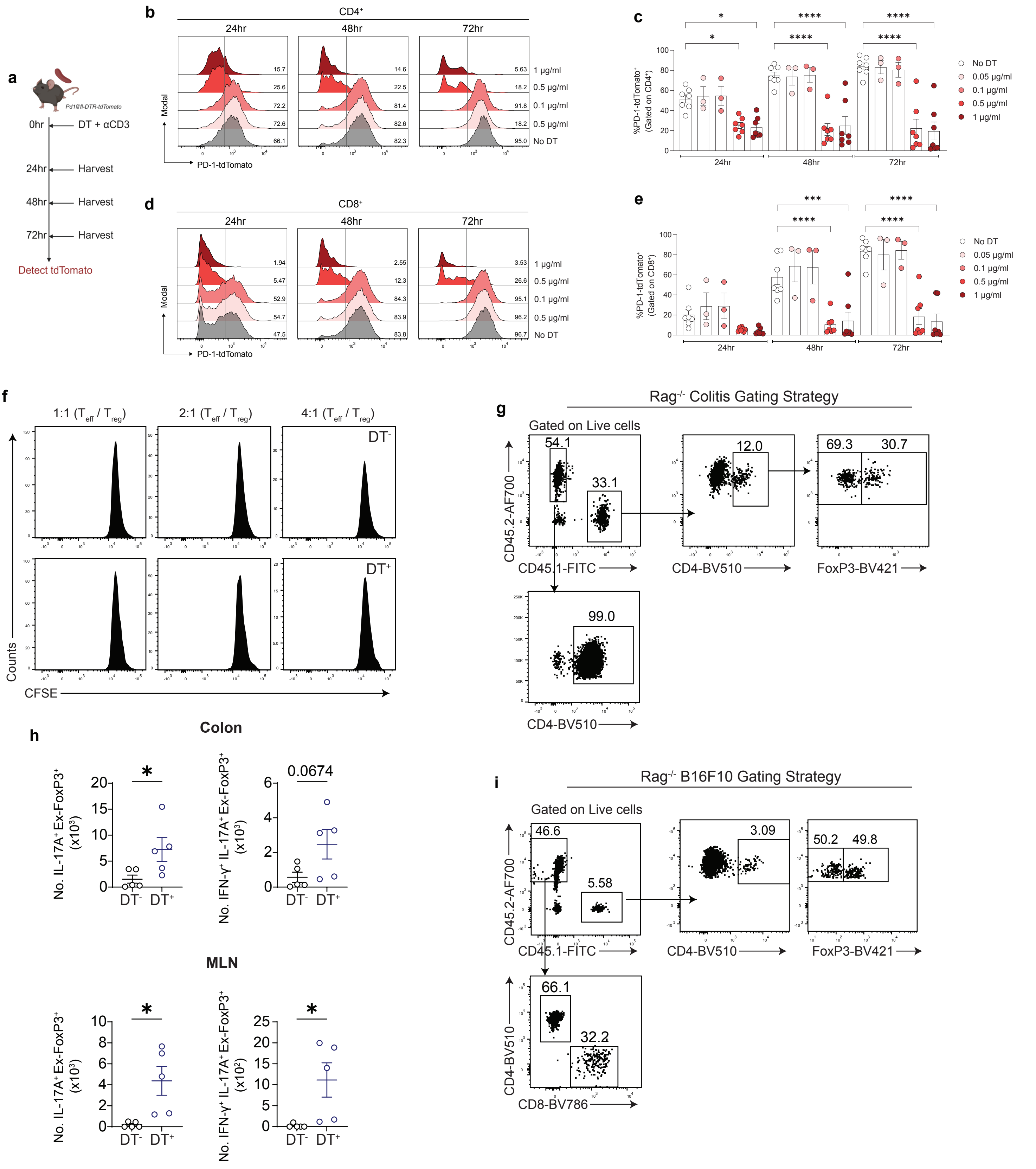


**Extended Figure 9**

### Figure 6

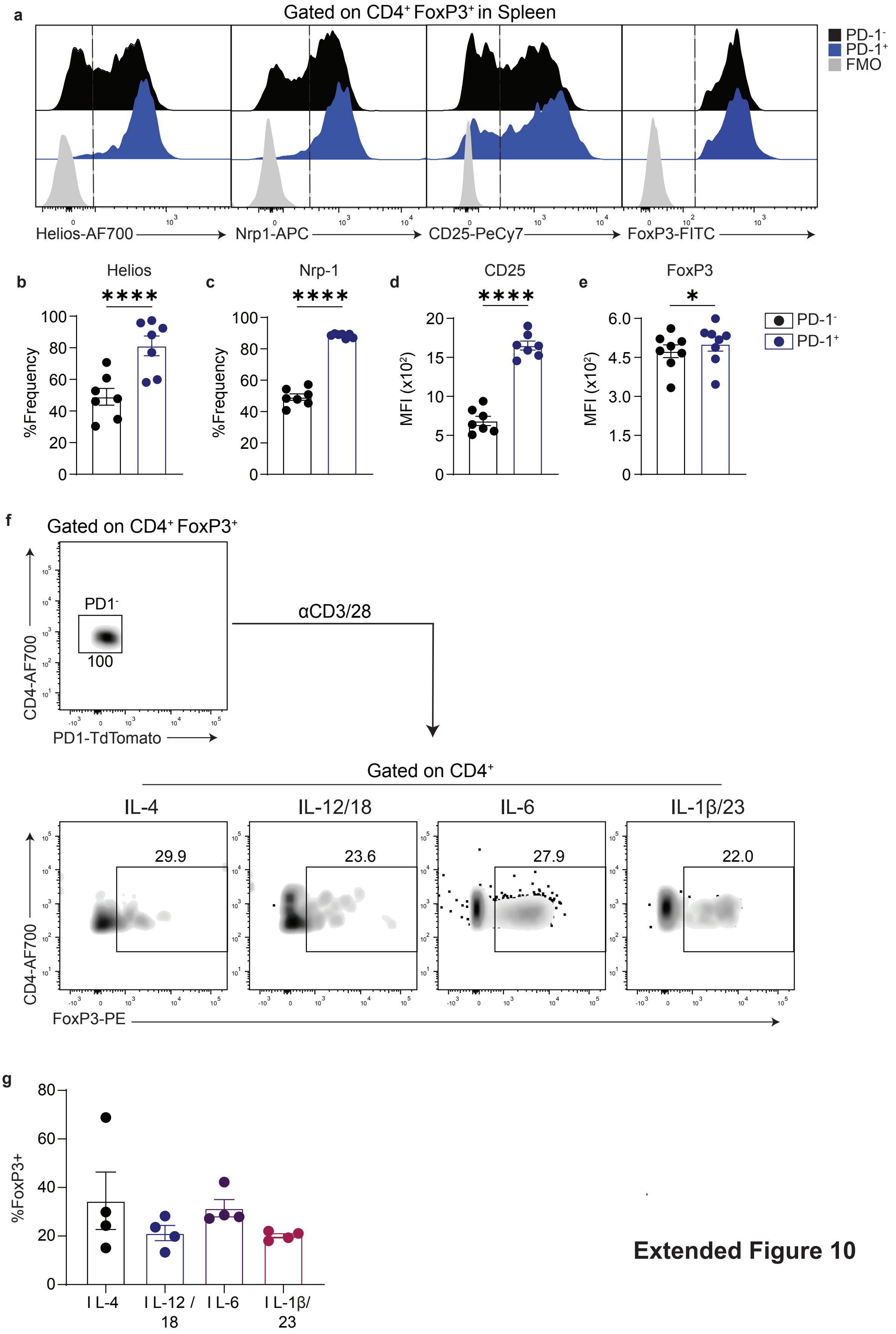


**Extended Figure 10**
